# HOXA9 forms a repressive complex with nuclear matrix-associated protein SAFB to maintain acute myeloid leukemia

**DOI:** 10.1101/2022.10.12.511919

**Authors:** Shuchi Agrawal Singh, Jaana Bagri, George Giotopoulos, Dhoyazan Azazi, Shubha Anand, Anne-Sophie Bach, Frances Stedham, Sarah J. Horton, Robin Antrobus, Jack W. Houghton, George S. Vassiliou, Daniel Sasca, Haiyang Yun, Anthony D. Whetton, Brian J.P. Huntly

## Abstract

*HOXA9* is commonly upregulated in acute myeloid leukemia (AML), where it confers poor prognosis. Characterising the protein interactome of endogenous HOXA9 in human AML, we identified a chromatin complex of HOXA9 with the nuclear matrix attachment protein-SAFB. *SAFB* perturbation phenocopied *HOXA9* knockout to decrease AML proliferation, increase differentiation and apoptosis *in vitro* and prolonged survival *in vivo*. Integrated genomic, transcriptomic and proteomic analyses further demonstrated that the HOXA9-SAFB-chromatin complex associates with NuRD and HP1γ to repress the expression of factors associated with differentiation and apoptosis, including *NOTCH, CEBP*δ, *S100A8*, and *CDKN1A*. Chemical or genetic perturbation of NuRD and HP1γ catalytic activity also triggered differentiation, apoptosis and the induction of these tumor-suppressive genes. Importantly, this mechanism is operative in other HOXA9-dependent AML genotypes. This mechanistic insight demonstrates active *HOXA9*-dependent differentiation block as a potent mechanism of disease maintenance in AML, that may be amenable to therapeutic intervention via therapies targeting the HOXA9/SAFB interface and/or NuRD and HP1γ activity.

**Key Points:**

- Identification of the endogenous human HOXA9 protein interactome in AML
- HOXA9 forms a repressive complex with S/MAR binding protein (SAFB) that is critical for the maintenance of AML by facilitating proliferation and preventing differentiation and cell death.
- The HOXA9/SAFB (H9SB) complex represses gene expression via recruitment of NuRD and HP1γ.

## Introduction

Acute Myeloid Leukemia (AML) is an aggressive blood cancer characterised by increased proliferation and self-renewal of immature cells of the myeloid series and a failure of differentiation within this compartment ^1^. A cardinal feature of AML is aberrant transcription and mutations that alter transcription factors, epigenetic regulators and genome structural proteins are frequent and recurrent ^2^. A common mediator of transformation downstream of many of these mutations, and overexpressed in ~70% of cases of AML, is the clustered homeobox transcription factor *HOXA9* ^3–5^. Overexpression of *HOXA9* also identifies a very poor-risk AML subgroup, with expression levels being highly predictive of prognosis ^6,7^.

However, the mechanisms whereby HOXA9 mediates transformation in the hematopoietic Stem and Progenitor Cell (HSPC) compartment remain relatively poorly understood. Previous studies have established its role as a transcriptional activator when associated with TALE homeobox proteins MEIS1 and PBX1/3 ^8,9^ more recent studies have shown this complex to drive proliferation and survival of leukemic cells through its interaction with multiple signalling pathways ^10–13^ and via stimulation of enhancer activation/modifications ^14,15^. Interestingly, *HOXA9* has been described to also have repressive function ^16^. However, the contribution of HOXA9-mediated repression to AML induction and maintenance remains unknown. Historically, studies of HOXA9 function have been somewhat hampered by the lack of reliable tools, with the majority of genomic and proteomic studies based on exogenous overexpression of tagged versions of HOXA9. Here, we make use of a commercial HOXA9 antibody validated in ChIP-seq ^10^ and, for the first time, we describe the protein interactome of endogenous HOXA9 in human AML cells. We then use these data to identify a novel HOXA9-repressive complex and characterize its functional and mechanistic role in AML maintenance.

## Methods

### Cell Lines

MOLM13, MV411, HL60 cells were cultured in RPMI1640 medium containing 10%FBS, 1%L-Glutamine and 1%PenStrep. OCIAML3 cells were cultured in MEMa medium containing 20%FBS, 1%L-Glutamine and 1%PenStrep. For treatment with inhibitors Panobinostat-LBH589 (Cat # S1030, Selleckchem) and Chaetocin (Cat # HY-N2019, MedChemExpress), cells were seeded in 6 well plate at density of 5×10^5^/ml in 3 ml media. Inhibitors were diluted to working stock of 10mM with DMSO and were used at final concentration of 4nM Panobinostat and 40nM Chaetocin. Cells were harvested for assays at specific time points.

### Rapid immunoprecipitation mass spectrometry of endogenous proteins (RIME)

RIME was performed as described earlier (Mohammed et al., 2016).

### CUTnRUN

CUTnRUN was performed using CUTANA CUT&RUN kit (Epicypher) following manufacturer’s instructions. 100,000 MOLM13 cells were used for the assay for each antibody. Antibodies are listed in the table. For Input, 100,000 cells were used and treated with MNase similar to antibody containing sample and processed as other samples. Libraries were prepared using 2ng of DNA using NEBNext UltraII DNA library preparation kit (Cat# E7645S, Illumina) and Dual index primer-Multiplex Oligos (Cat# E7600S, Illumina).

### Cloning shRNAs into lentiviral vector (pLKO-TetOn)

shRNA against human HOXA9 and SAFB were cloned into pLKO “all-in-one” system for the inducible shRNA expression as described earlier ^17,18^.

Detailed methods are provided in supplementary section materials and methods.

## Results

### A proteomic screen identifies HOXA9 to interact with matrix (S/MAR) binding proteins

In order to define the HOXA9-interacting proteome in acute leukemia, we performed immunoprecipitation of endogenous HOXA9 (HOXA9-IP) followed by mass spectrometry in the HOXA9-dependent (Faber et al., 2009) *MLL-AF9*-rearranged MOLM13 AML cell line. Two replicates of immunoprecipitated samples were analysed by label free quantification mass spectrometry (LFQMS), giving a high degree of concordance. (Supplemental Figure 1A). There were 324 significantly enriched proteins (>2 log-fold in HOXA9 vs IgG) (Figure 1A, Supplemental Figure 1B, supplemental table 1). Consistent with its transcriptional role, chromatin binding proteins coimmunoprecipitated with HOXA9. These included factors involved in transcription initiation, mRNA processing, basal transcription factors, acetyltransferases and lineage specific transcription factors including CEBPα (Figure 1B). Interestingly, the scaffold/matrix attachment region (S/MAR) proteins SATB1, SATB2, and SAFB were also greatly enriched. However, while SATB1 has been described to be rearranged in AML ^20^ and appears to regulate the tumour suppressor function of PU.1 in AML ^21^ and the repressive function of SATB2 is required to block B-cell differentiation in BCR-ABL+ B-ALL ^22^, there are no previous reports of a role for SAFB in leukemia. To gain additional insights into the role of these S/MAR proteins in leukemia, we analyzed our previously published CRISPR-dropout data from a screen of five AML cell lines ^23^. This demonstrated that depletion of *SAFB* significantly inhibited the growth of all tested AML cell lines, while depletion of *SATB1* and *SATB2* did not (Figure 1C). These findings were corroborated using another, independent CRISPR-dropout screen encompassing 14 human AML cell lines (Supplemental Figure 1C) ^24^, suggesting a novel, essential function of *SAFB* for the growth of AML cells.

**Figure 1.**
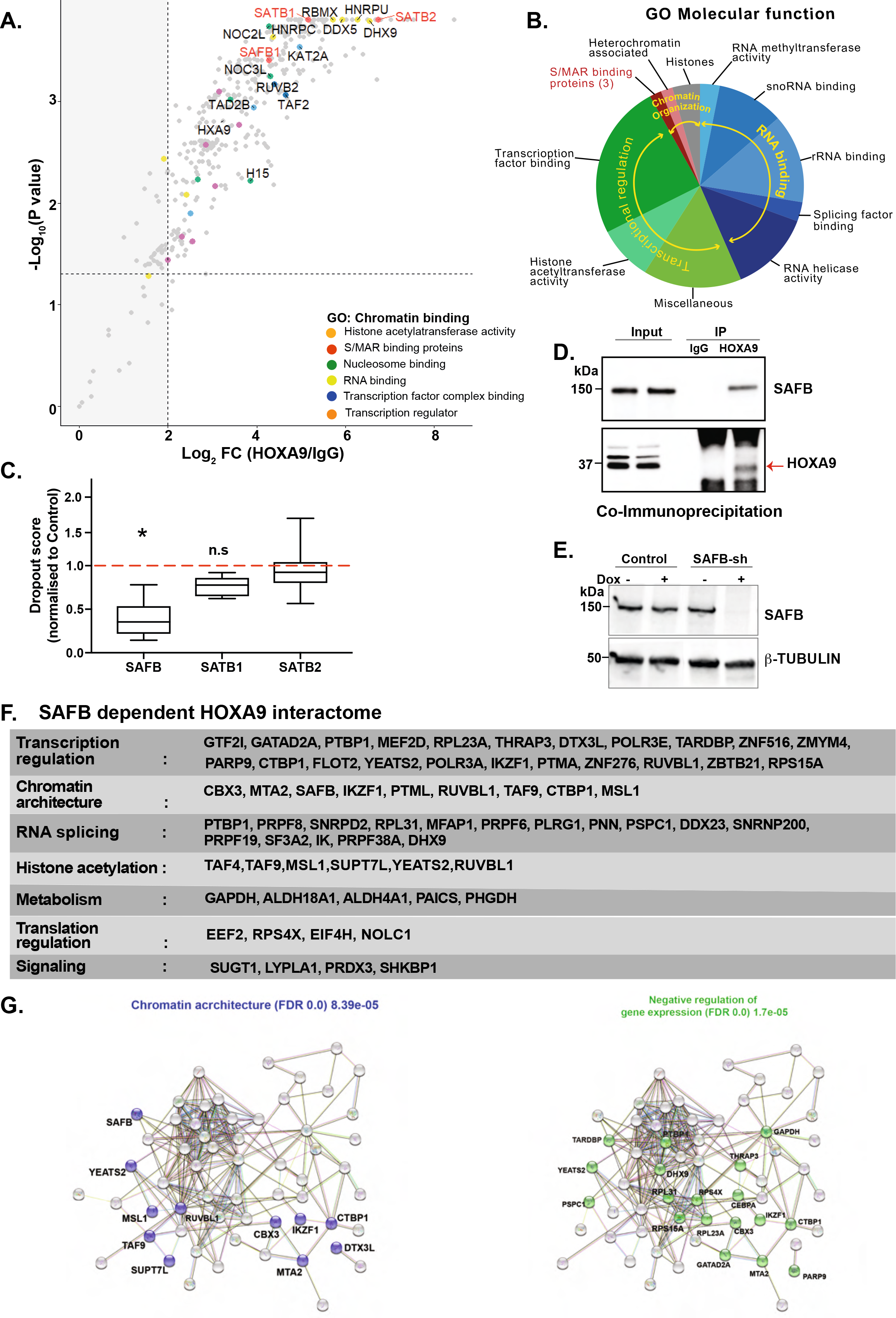
HOXA9 interacts with matrix binding (S/MAR) protein SAFB. A) Volcano plot displaying label free MS quantification of HOXA9 pull down in MOLM13 cells. The plot shows log2 ratios of averaged peptide MS intensities between HOXA9-IP and control-IP (IgG) eluate samples (X axis) plotted against the negative log10 p values (Y axis) calculated across the replicate data sets (one tailed Student’s t test, n=2 replicates). Maximum upper values were set for the X and Y axis to accommodate all detected proteins in the plot. A dashed horizontal line marks p value = 0.05. Vertical dashed line marks enrichment >2 log2 fold. Chromatin binding proteins, pulled-down by HOXA9 are coloured as indicated, and selected protein names are shown. The full dataset is given in Table S1. B) Summary of proteins pulled-down by HOXA9-IP, categorised according to molecular function from gene ontology analyses, FDR <0.000001. C) Box plot represents depletion of *SAFB, SATB1, SATB2* by CRISPR in 5 AML cell lines. Dropout score was calculated by normalising to control cells (non-AML) (Tzelepis *et al*., 2016). D) Western blot analyses validating the HOXA9 and SAFB interaction in MOLM13 cells via co-immunoprecipitation. E) Representative western blot showing SAFB knockdown via doxycycline inducible shRNAs in MOLM13 cells at 48hr post-induction. β-Tubulin used as loading control. F) Summary of SAFB-dependent HOXA9 interacting proteins (n=65), FDR<0.05, log2FC >1.5. G) String network analysis demonstrating HOXA9 interacting proteins (SAFB-dependent) involved in chromatin architecture, (left panel, blue) or involved in negative regulation of gene expression (right panel, green).

We next confirmed the SAFB interaction with HOXA9 in MOLM13 cells by co-immunoprecipitation (Figure 1D). To determine whether SAFB is required for the identified HOXA9 chromatin interactions, we performed Rapid Immunoprecipitation Mass spectrometry of Endogenous proteins (RIME)^25^ for HOXA9 in MOLM13 cells, comparing SAFB knockdown to control (Figure 1E). We identified 65 specific proteins that interacted with HOXA9 only in the presence of SAFB (Figure 1F). Of note, proteins involved in chromatin architecture and gene repression were prominent within this SAFB-dependent subset (Figure 1G).

### SAFB phenocopies HOXA9 to drive proliferation and prevent differentiation and apoptosis in leukemic cells

Our previous global CRISPR-screen data suggested that SAFB has a functional requirement across multiple AML genotypes. Therefore, to specifically dissect the role of SAFB in HOXA9-dependent AML, we used single guides to individually target *SAFB* and *HOXA9* expression in MOLM13 cells ^26^. *HOXA9* depletion led to reduced cell growth compared to control cells (Figure 2A-B) and an increase in myeloid differentiation and apoptosis (Figure 2C-D). Of note, *SAFB* depletion fully phenocopied HOXA9 depletion, suggesting a mechanistic interaction between the associating proteins.

**Figure 2.**
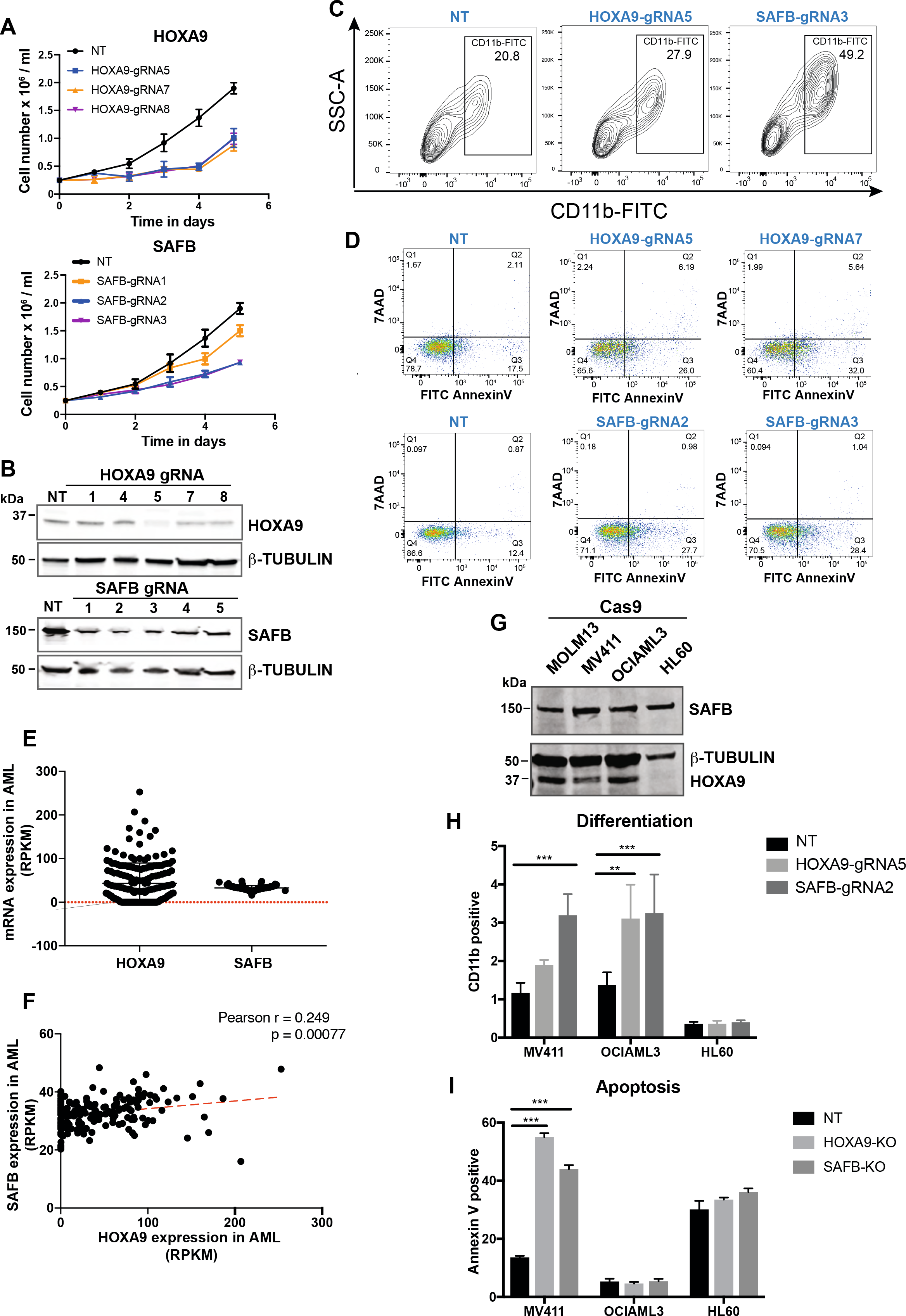
SAFB phenocopies HOXA9 in leukemic cells. A) Growth kinetics of MOLM13-Cas9 cells transduced with gRNA targeting *HOXA9* or *SAFB*. The data are shown as average of biological replicates (n=3) ± SD. 2way ANOVA test, * p<0.01. NT=Non-targeting control gRNA. B) Western blot analyses showing the knockdown efficiency of HOXA9 and SAFB gRNAs in MOLM13 cells. C) Flow-cytometric analyses of CD11b surface expression in MOLM13-cas9 cells transduced with gRNA targeting *HOXA9* or *SAFB*. Cells were stained for CD11b-FITC, after 3 days of transduction. Contour plots shown here are representative of 3 independent biological replicates. NT=Non-targeting control gRNA. D) Apoptosis in MOLM13-cas9 cells transduced with gRNA targeting *HOXA9* or *SAFB* five days post-transduction as measured by AnnexinV and 7AAD staining. Plots are representative of 3 independent biological experiments. NT=Non-targeting control gRNA. E) The mRNA expression of *HOXA9* and *SAFB* in human AML primary samples (n=179), AML-TCGA data set. F) Correlation between *HOXA9* and *SAFB* mRNA expression in human AML primary samples (n=179), AML-TCGA data set. G) Western blot analyses showing expression of SAFB and HOXA9 in AML cells lines. β-TUBULIN loading control. H) CD11b surface expression in AML cell lines, 3 days post-transduction with gRNA targeting *HOXA9* or *SAFB* (mean± SD, n=3). Statistical significance calculated using 2way ANOVA test, ** p<0.01. NT=Non-targeting control gRNA. I) Apoptosis measured by Annexin V positivity in AML cell lines, 3 days after transduction with gRNA targeting *HOXA9* or *SAFB* (mean ± SD, n=3). Statistical significance was calculated using 2way ANOVA test, ***| p<0.01. NT=Non-targeting control gRNA.

To further assess its role in primary AML, we next assessed *SAFB* expression using whole transcriptome data from 179 primary AMLs (TCGA Research Network https://www.cancer.gov/tcga) across all major subtypes. *SAFB* mRNA levels were remarkably homogeneous across multiple AML samples/subtypes (Figure 2E) but a statistically significant positive correlation with *HOXA9* expression was observed, even though *HOXA9* levels are more heterogenous across AML (Figure 2F). To further test the functional interaction, we targeted *SAFB* expression by CRISPR-Cas9 in two additional *HOXA9*-dependent AML cell lines (MV411- *MLL-AF4* rearranged and OCIAML3 – *NPM1*-mutated), as well as a *HOXA9*-independent (HL60) AML cell line (Figure 2G). Similar to the *MLL-AF9+* MOLM 13 cell line, we could demonstrate that depletion of *SAFB* induced differentiation of *HOXA9*-dependent cell lines (MV411 and OCIAML3), but had no impact on HL60 cells (Figure 2H). The induction of apoptosis was also observed in MV411, whereas no apoptosis was observed in OCIAML3 and HL60 cells (Figure 2I).

### HOXA9 and SAFB drive leukemic growth *in vivo*

To determine whether SAFB and HOXA9 are required for leukemia maintenance *in vivo*, we utilised the temporal control of a doxycycline inducible shRNA system (pLKO) to induce knockdown only after disease establishment had been demonstrated. We first confirmed the functional effects of *SAFB* and *HOXA9* knockdown on MOLM13 cell growth *in vitro*. *SAFB* and *HOXA9* knockdown in the presence of doxycycline (Figure 3B) phenocopied CRISPR-editing to substantially reduce the growth of MOLM13 cells (Figure 3A), reduce the proportion of cells in S-phase (Supplemental Figure 2A), induced myeloid differentiation followed by apoptosis and reduced methylcellulose colony formation (Figure 3C-3E and Supplemental Figure S2B). We next injected NSG mice with MOLM13-sh cells that expressed a luciferase reporter-gene in addition to an inducible shRNA vector to knockdown either *SAFB* or *HOXA9* expression (experimental scheme Figure 3F). Three days post injection, engraftment and disease induction were confirmed by bioluminescence imaging and the baseline was calculated for all 3 cohorts of mice (Figure 3G). In accordance with our *in vitro* experiments, knockdown of *HOXA9* and *SAFB* significantly delayed disease progression and prolonged the survival of mice compared to the control cells (Figure 3H-J). Together, these data confirm a functional as well as a physical interplay between the two proteins, with SAFB phenocopying HOXA9 function to support leukemic growth and prevent differentiation of leukemic cells both *in vitro* and *in vivo*.

**Figure 3.**
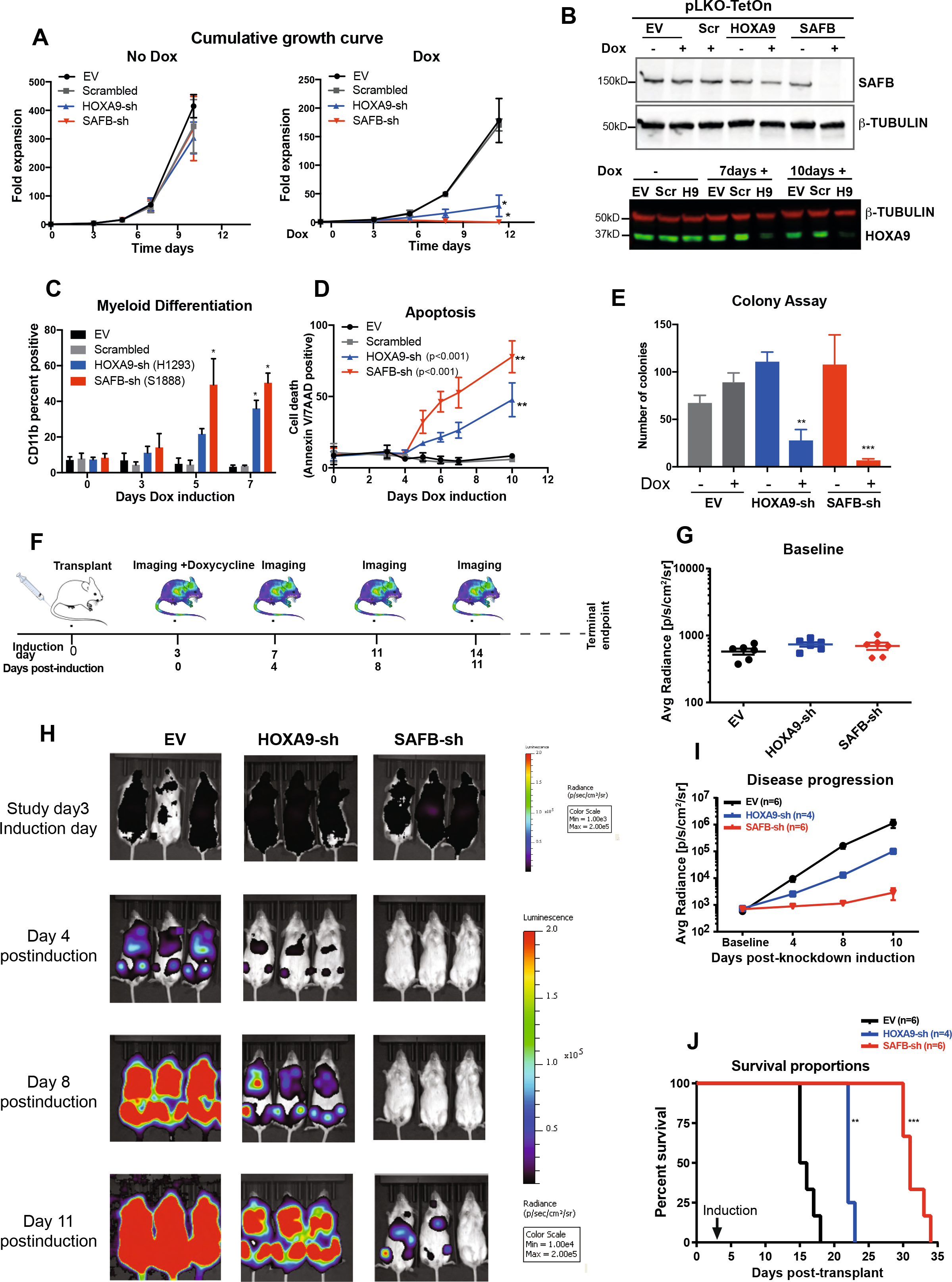
HOXA9 and SAFB drive leukemic growth *In vivo*. A) Cumulative growth of MOLM13 cells, lentivirally expressing shRNA against *HOXA9* (blue), *SAFB* (red), scrambled sequence (grey), empty vector (EV, black). The data shown as average of biological replicates (n=3) ± SD. 2way ANOVA test, * p<0.01 B) Western blot analyses showing knockdown efficiency of SAFB shRNAs after 48hr post-induction with doxycycline in MOLM13 cells (upper panel). β-TUBULIN used as loading control. Lower panel demonstrates the HOXA9 knockdown after 7 and 10 days of doxycycline induction in MOLM13 cells. EV=empty vector, Scr=Scrambled, H9=HOXA9sh C) Myeloid differentiation accessed by CD11b surface expression by flow cytometry 5 days post-induction with doxycycline. Data shown as average of biological replicates (n=4) ± SD. Statistical significance calculated using 2way ANOVA test, *p<0.01 D) Percentage of Annexin V and 7AAD positive MOLM13 cells 5 days after doxycycline treatment (1.5μg/ml). (Mean± SD, n=4}. 2way ANOVA test, **p<0.001. E) Colony forming assay of MOLM13 cells expressing *HOXA9* or *SAFB* shRNA in the presence or absence of doxycycline (1.5μg/ml). Bar graph shows the average value of 3 independent experiments +SD, 2way ANOVA test **p<0.001, ***p<0.0001. F) Schematic of xenotransplant experimental design. G) Bioluminescent radiance 3 days post-injection and prior to sh-RNA induction (baseline), in all 3 cohorts in all animals. H) Serial bioluminescence imaging of mice transplanted with luciferase-labelled shRNA-expressing MOLM13 cells at indicated time points. I) Bioluminescence at indicated time points shows disease progression over time. J) Kaplan Meir plot showing the survival of mice transplanted with MOLM13 cells expressing indicated shRNA. Log Rank test was performed (**p<0.01, ***p<0.001).

### SAFB co-localizes with HOXA9 genome-wide

We next probed the function of the putative HOXA9/SAFB complex. To detail HOXA9/SAFB localisation on chromatin, we performed CUT&RUN ^27^ for HOXA9 or SAFB in MOLM13 cells. The HOXA9 antibody utilised has been previously shown to enrich endogenous HOXA9 in ChIP-seq experiments ^10^ and both HOXA9 or SAFB antibodies gave consistent results and compared favourably with ChIP-Seq (Supplemental Figures 3A-C). We identified high confidence binding sites for HOXA9 (n=39777) and for SAFB (n=12672) in MOLM13 cells (Supplemental Figure 3B). Of note, we found that a subset of HOXA9 peaks specifically co-localize with the majority of SAFB peaks (n=10,262, Figure 4A-B and Supplemental Figure 3D, supplemental table2). Among those, one third of HOXA9-SAFB co-bound peaks were found at promoters (n=3845) with the remainder as distal peaks (n=6417). Motif analyses of HOXA9/SAFB co-occupied loci showed strong enrichment for hematopoietic transcription factor binding sites, including PU.1 (ETS), RUNX1and CEBPA (Supplemental Figure 3E).

**Figure 4.**
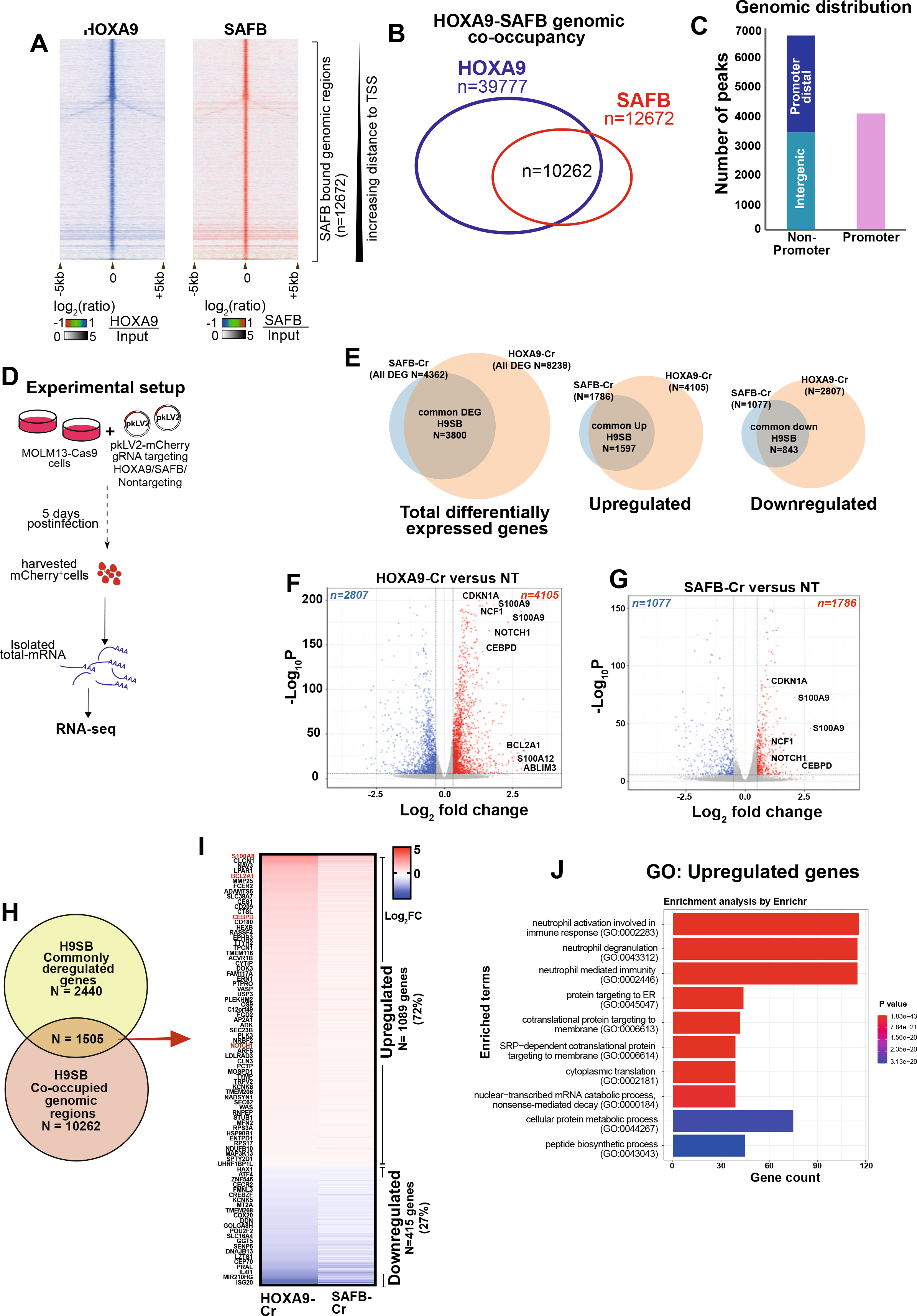
HOXA9-SAFB together repress the transcription of myeloid differentiation genes. A) Heat maps of HOXA9 or SAFB signal (relative to Input) from CUT&RUN experiments in MOLM13 cells. The Y-axis represents individual regions centred at SAFB bound genomic peaks (±5kb). Regions were sorted according to increasing distance to TSS. The relationship between coloring and signal intensity is shown in the bar at the bottom of the plot. B) Schematic representation of overlapping regions occupied by HOXA9 and SAFB. C) A bar graph showing the genomic distribution of HOXA9-SAFB co-bound regions. D) Schematic of experimental setup for RNA sequencing using MOLM13-cas9 cells transduced with *HOXA9* or *SAFB* gRNA. E) Venn diagram shows overlap between total differentially regulated genes in *HOXA9-* and *SAFB*-CRISPR knockout cells. Data analysed using three independent replicates at p<0.001; for upregulated or downregulated genes threshold set to 1±0.2. Hypergeometric test p value <0.00001 for all overlaps. F) RNA-sequencing volcano plot showing genes downregulated (left, blue) and genes upregulated (right, red) in *HOXA9-*CRISPR compared to non-targeting control MOLM13 cells (n=3 in both conditions). G) RNA-seq in *SAFB-*CRISPR MOLM13 cells as described in F. H) Venn diagram shows the overlap between genes commonly dysregulated upon *HOXA9* and *SAFB* perturbation in MOLM13 cells (yellow) and genes that are linked to HOXA9-SAFB co-occupied genomic regions (orange). List of these genes provided in Table S4. Hypergeometric test; p value 2.438e-13. I) Heatmap showing expression Log2 fold change in (HOXA9-Cr Vs NT) or (*SAFB-*Cr vs NT) MOLM13 cells of those 1505 target genes. The relationship between colouring and expression value is shown in the bar to the right side of the plot. J) Gene ontology analyses for genes that demonstrate occupancy of HOXA9-SAFB (within 50kb to TSS), and are upregulated in response to HOXA9 or SAFB perturbation in MOLM13 cells. Gene count is plotted on X-axis, with the significance p-value given by the color code on the bar to the right.

As expected by their SAFB binding, these co-bound loci demonstrated characteristic sequence features of S/MAR regions, such as AT-proportion, topoisomerase II binding, kinked or curved DNA ^28,29^ (Supplemental Figure 4A). Of note, co-binding could be demonstrated at or near the classic S/MAR regions of the *MYC* ^30^, *TOPI, TOP2A* ^31^ and β*-GLOBIN* ^32^ genes (Supplemental Figure 4B). Taken together these analyses demonstrate that a significant subset of the HOXA9 cistrome associates on chromatin with the majority of SAFB-bound loci at S/MAR-associated cis-regulatory regions.

### Transcriptional consequence of HOXA9/SAFB depletion in leukemic cells

We next profiled the transcriptome of *HOXA9* and *SAFB* depleted MOLM13 leukemic cells using RNA seq (Figure 4D). Differential gene expression analyses demonstrated a total of 8238 altered genes in *HOXA9*-depleted cells, whereas *SAFB*-depletion altered the expression of 4362 genes (p<0.01). The majority of genes affected by SAFB-depletion (87%) were also altered by HOXA9-depletion (Figure 4E). Furthermore, gene set enrichment analyses (GSEA) of differentially regulated genes following either *HOXA9* and *SAFB* depletion enriched for a HOXA9 signature (Supplemental Figure 5A). Within the overlapping genes, differential expression was observed in both directions (threshold set to 1±0.2 to measure any significant change in the expression, overlap among up-regulated or down-regulated genes n=89%, or n=78% respectively, p<0.01) (Figure 4E-G, Supplemental Table 3). Gene ontology (GO) enrichment analyses demonstrated marked enrichment for terms associated with myeloid differentiation among commonly up-regulated genes, whereas, interferon and viral responsive genes were negatively affected by the loss of *HOXA9* and *SAFB* (Supplemental Figure 5B).

To identify direct targets of HOXA9 and SAFB, we integrated HOXA9/SAFB CUT&RUN-seq data with RNA-seq following *HOXA9/SAFB* perturbation. Of interest, 61% of the commonly dysregulated genes (n=1505/2440), perturbed both by loss of *HOXA9* and *SAFB*, demonstrated a HOXA9/SAFB binding peak within 50kb of their TSS (Figure 4H, Supplemental Table 4), with the majority (72%) up-regulated following *HOXA9/SAFB* perturbation, in keeping with our developing hypothesis of a repressive role for the putative HOXA9/SAFB (H9SB) complex (Figure 4I). GO analyses revealed genes associated with myeloid differentiation and function to be significantly enriched in this up-regulated gene set (including *S100A8, S100A9, NOTCH1, CEBP*δ, *CDKN1A, BCL2A1, S100A12, SERPINA1*), whereas downregulated genes were enriched in interferon response pathways (Figure 4J and Supplemental Figure 5C).

### NOTCH and CEBPδ are targets of the HOXA9/SAFB (H9SB) repressive complex

The *NOTCH1* and *CEBP*δ genes were de-repressed upon HOXA9 or SAFB perturbation and co-occupancy of HOXA9-SAFB was observed proximal to their loci (Figure 5A and F). Similar relationships with HOXA9/SAFB binding and up-regulated gene expression upon their perturbation were also observed for the *CDKN1A*, *S100A8, S100A9 S100A12, BCL2A1* genes (Supplemental Figure 6A).

**Figure 5.**
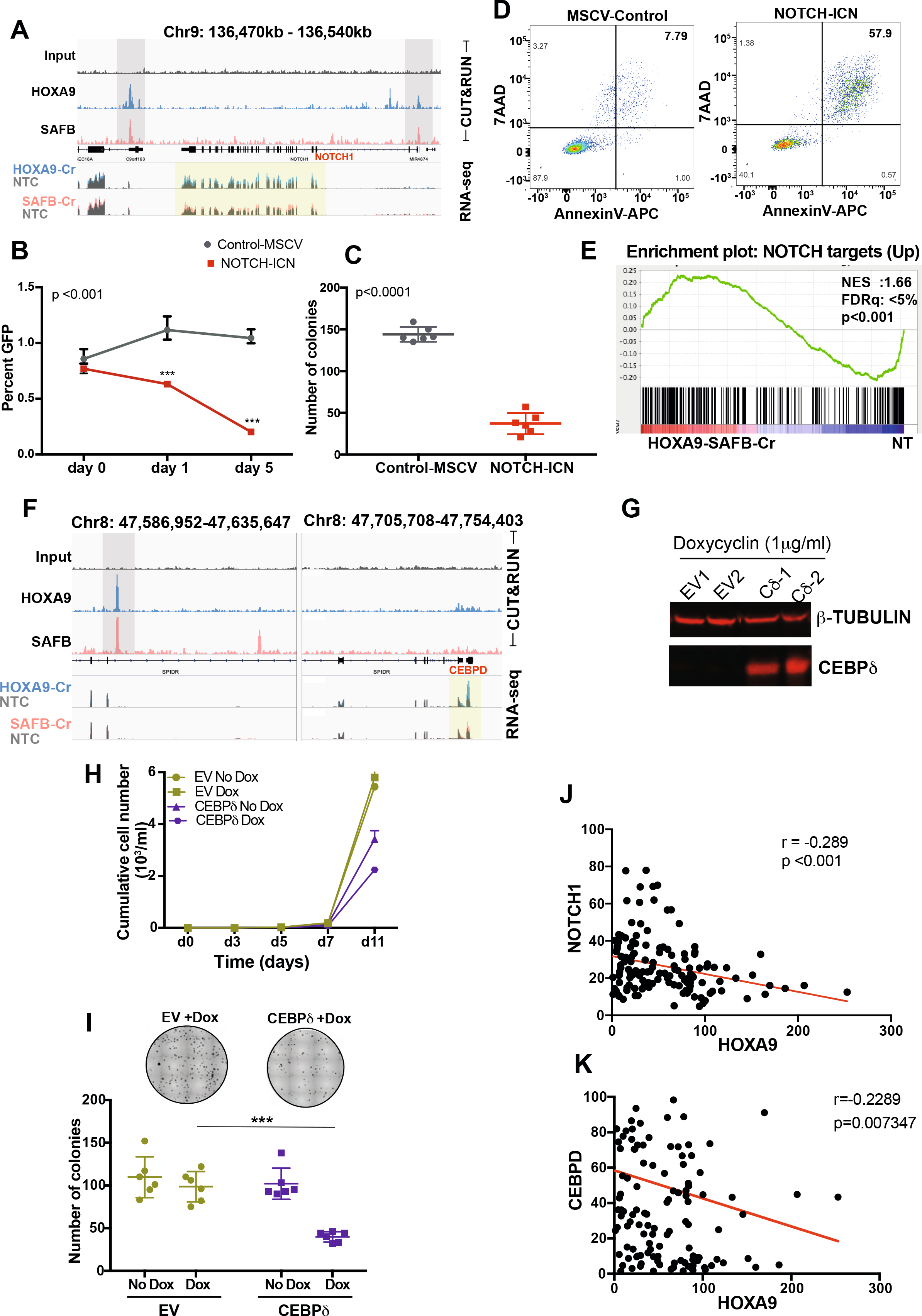
NOTCH and CEBPδ are targets of the HOXA9/SAFB (H9SB) repressive complex. A) Genome browser tracks demonstrate the enrichment of HOXA9 (blue) and SAFB (red) at the representative *NOTCH1* locus in Hg38 genome obtained from CUT&RUN sequencing in MOLM13 cells. Lower two tracks show the RNA seq data, expression of *NOTCH1* to be upregulated in *HOXA9*-Cr (overlain blue) and *SAFB*-Cr (overlain red) compared to control-non targeting (NT) (grey). B) Competition assay between GFP positive NOTCH-ICN or MSCV-control vector-expressing MOLM13 cells. Equal numbers of GFP positive cells were seeded at day0 and the GFP percentage was measured 3 and 5 days after by flowcytometry. Data shown as average of 3 biological replicates ± SD. C) Colony-forming assay of GFP positive MOLM13-NOTCH-ICN or MOLM13-MSCV-control cells. Bar graph shows the average value of 3 independent experiments ± SD. D) Apoptosis in MOLM13-NOTCH-ICN or MOLM13-MSCV-control cells measured by AnnexinV and 7AAD positive cells by flow-cytometry. Plots are representative of 3 biological independent replicates. E) GSEA for HOXA9-SAFB commonly regulated genes (n=2440) enriches for NOTCH signature. F) Genome browser track representation of another exemplar locus, *CEBPδ* description as A. G) Western blot shows the expression of CEBPδ in MOLM13 cells upon doxycycline treatment for 2 days. β-TUBULIN loading control. H) Cumulative cell growth of MOLM13 cells in CEBPδ overexpressing cells +doxycycline. The experiment shown here is average of 3 independent replicates obtained from two independent clones. Error bars represent ± SD. I) Colony forming assay of MOLM13-CEBPδ or MOLM13-control cells ±doxycycline. Upper panel - photomicrographs showing representative plates. Lower panel - scatter plot shows 3 independent experiments in duplicates, showing median ± SD. Significance calculated using one way-ANOVA, p<0.001. J) Correlation between *HOXA9* and *NOTCH1* mRNA expression in human AML primary samples (n=165, TCGA data set). K) Correlation between HOXA9 and CEBPδ mRNA expression in human AML primary samples (n=165, TCGA data set).

We next sought to link de-repression of specific genes and activation of the differentiation programme with abrogation of the leukemia maintenance phenotype. Upon retroviral expression of an activated form of NOTCH (NOTCH-ICN) in MOLM13 cells, we saw a strong impairment of growth/proliferation and clonogenic potential, accompanied by a marked increase in apoptosis (Figure 5B-D). Furthermore, GSEA analyses on a pre-ranked list of differentially expressed genes from *HOXA9/SAFB* depleted MOLM13 cells demonstrated a striking correlation with NOTCH signatures (Figure 5E). To further extend our paradigm, we also assessed the effects of re-expressing *CEBP*δ in MOLM13 cells using an inducible system (Figure 5G). Similarly, *CEBP*δ expression attenuated MOLM13 cell growth and inhibited clonogenic growth in semi-solid methylcellulose medium (Figure 5H-I). Moreover, analyses of 179 primary AML patients across all major subtypes revealed a negative correlation between *NOTCH1 or CEBP*δ and *HOXA9* mRNA expression level (Figure 5J-K, Supplemental Figure 6B).

Taken together, these data demonstrate that, in concert with SAFB, HOXA9 mediates repression of *NOTCH, CEBP*δ and other myeloid differentiation-associated genes to actively maintain the differentiation block associated with AML.

### HOXA9/SAFB forms a repressive complex on chromatin with NuRD and HP1γ

We next sought to explore the mechanism of gene repression at chromatin mediated by the H9SB complex. We therefore performed further proteomic analyses to identify chromatin-associated protein interactors of the complex using RIME (Figure 6A) ^25^. Endogenous SAFB or HOXA9 were precipitated from MOLM13 nuclei after formaldehyde crosslinking and compared with IgG (Supplemental Figure 7).

**Figure 6.**
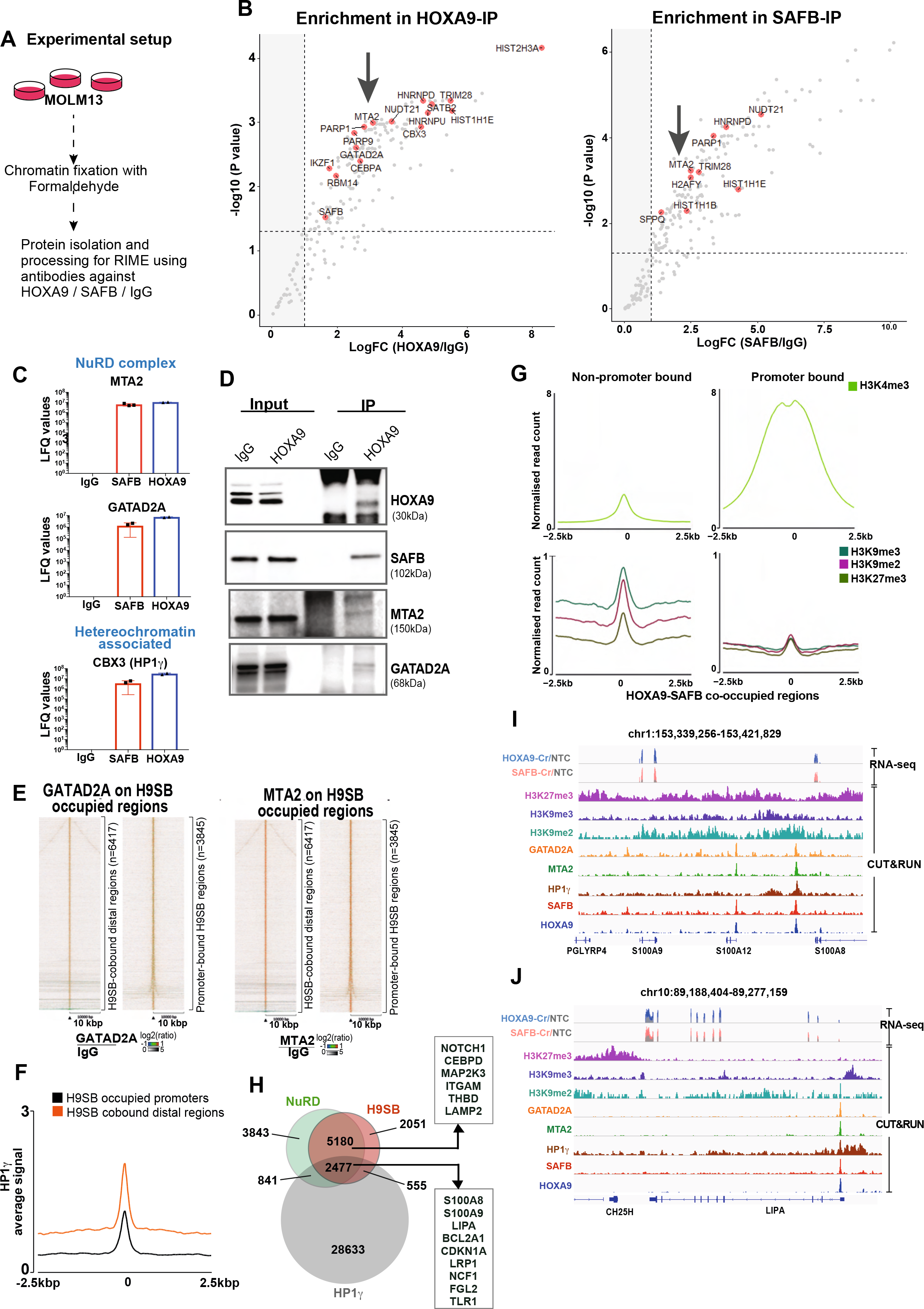
H9SB forms a repressive complex on chromatin with NuRD and HP1γ. A) Schematic of experimental setup for proteomic RIME screen using MOLM13 cells. B) Volcano plot displaying the label free quantitative MS result of HOXA9 and SAFB immunoprecipitation in MOLM13 cells. The plot shows log2 ratio of averaged peptide MS intensities between HOXA9-IP Vs IgG (left panel) or SAFB IP Vs IgG (right panel) samples (X-axis), plotted against the negative log10 P values (Y axis) calculated across the triplicate datasets. (Student t test, n=3 technical replicates). Dashed black line marks 1.5 log FC. Chromatin associated proteins enriched in HOXA9 or in SAFB pull-downs are marked as red dots. The complete list of SAFB-HOXA9 commonly enriched proteins is given in Table S1. C) Bar graph displays the enrichment of NuRD complex members (MTA2 and GATAD2A) or heterochromatin protein HP1γ in HOXA9 or SAFB immunoprecipitated samples. The data shown here are intensities from label free quantification values from mass spectrometric analyses from all replicates for IgG (n=3), SAFB (n=3), HOXA9 (n=2) pull downs, average ±SD. D) Western blots showing HOXA9 interaction with SAFB, MTA2 and GATAD2A by co-immunoprecipitation of endogenous HOXA9-pulldown in MOLM13 cells. E) Heat maps of MTA2 and GATAD2A signal (relative to IgG) on HOXA9-SAFB co-occupied genomic regions; promoter bound (left panel), distal bound (right panel) measured by CUT&RUN in MOLM13 cells. The Y-axis represents individual regions centred at HOXA9-SAFB bound genomic regions (±10kb). Regions were sorted according to increasing distance to TSS. The relationship between colouring and signal intensity is shown in the bar at the bottom of the plot. F) Average signal for HP1γ enrichment (intensity on Y-axis as normalised read count) centred at promoter or distal HOXA9-SAFB-co-occupied regions as measured by CUT&RUN in MOLM13 cells. G) Average signal of indicated histone modification (intensity on Y-axis as normalised read count) active-H3K4me3 (top panel) and repressive-H3K9me2, -H3K9me3 and -H3K27me3 (lower panel) centred at non-promoter or promoter bound HOXA9-SAFB-co-occupied regions determined by CUT&RUN in MOLM13 cells. H) Venn diagram showing the overlap of high-confident NuRD (GATAD2A+MTA2), and HP1γ peaks with H9SB co-bound genomic regions in MOLM13 cells. The differentiation-associated target-genes of the H9SB-repressive complex also upregulated upon HOXA9-SAFB perturbation are highlighted. I) and J Genome browser track shows the co-localization of NuRD complex, HP1γ and repressive modifications on HOXA9-SAFB enriched region in the Hg38 genome, obtained from CUT&RUN sequencing in MOLM13 cells.

Considering only high-confidence proteins present in all replicates, we identified 162 proteins significantly enriched in SAFB-RIME (Figure 6B, details are provided in the methods section). Overlapping this dataset with proteins enriched in HOXA9 RIME identified 79 proteins (the complete list of HOXA9/SAFB interacting proteins is provided in Supplemental Table 1). String network analyses [https://string-db.org/] functionally annotated these proteins ^33^ to be highly enriched for negative regulation of gene expression (47/79 proteins; FDR=1.33e-27). Clustering these proteins based on their molecular function resulted in 3 distinct clusters; RNA binding or splicing proteins (cluster 1), ribosomal proteins (cluster 2) and chromatin-associated proteins (cluster 3) (Supplemental Figure 8A). Cluster 3 included members of the NuRD (nucleosome remodelling and histone deacetylase) complex (MTA2 and GATAD2A) and heterochromatin protein CBX3 (also known as HP1γ) (Figure 6B-C, Supplemental Table 1). All three proteins showed strong enrichment compared to IgG controls using the MaxLFQ approach ^19^ (Figure 6C). Moreover, interaction of HOXA9 with MTA2 and HP1γ seemed to be SAFB dependent (Figure 1F and 1G). Further to this, we validated the physical association of HOXA9/SAFB with NuRD complex members by co-immunoprecipitation, implying the formation of a HOXA9-repressive complex in AML (Figure 6D).

To further gain insight into the genome wide interaction of the NuRD complex and HP1γ at HOXA9/SAFB bound loci, we performed CUT&RUN for GATAD2A, MTA2 and HP1γ in MOLM13 cells (Supplemental Figure 8B). These data further demonstrated that NuRD complex members (MTA2, GATAD2A) exhibited considerable genome-wide overlap with HOXA9/SAFB co-bound genomic regions (7657 overlapping regions out of 10262 (75%) H9SB co-bound regions) (Figure 6E, and 6H). This overlapping pattern was observed at similar frequencies for both promoter as well as for distal regions (Figure 6E). Although HP1γ enrichment was more modest, we did observe a clear signal in the proximity of HOXA9/SAFB bound genomic regions, with distal regions showing higher enrichment compared to promoter-bound regions (Figure 6F). Further analyses revealed that 32% (2477/7657) of the NuRD-H9SB overlapping regions also harboured HP1γ binding (Figure 6H, Supplemental Table 5). Moreover, perturbation of HOXA9 and SAFB commonly affected the expression of 600 genes (differentially expressed) annotated to these peaks (within 50kb), with 420 (70%) of these up-regulated genes. Of note, the up-regulated genes again demonstrated the association with myeloid differentiation in GO analyses (Figure 6H, and Supplemental Figure 9C).

Assessing histone modifications at the HOXA9/SAFB co-occupied loci revealed higher enrichment of repressive histone modifications (H3K27me3, H3K9me2, H3K9me3) compared with HOXA9-only occupied loci (Supplemental Figure 9A), with repressive histone modifications particularly enriched at non-promoter bound peaks (Figure 6G and Supplemental Figure 9B). However, these loci also exhibited enrichment for H3K4me3 although, as expected, enrichment was more prominent at promoter regions (Figure 6G, Supplemental Figure 9D),). Exemplar loci demonstrating co-occurrence of NuRD, HP1γ and repressive histone modifications with HOXA9-SAFB that correlated with de-repression of the associated genes upon HOXA9-SAFB perturbation are shown in Figures. 6I-J and Supplemental Figure 9D.

Having demonstrated the global co-binding of repressive complexes at HOXA9/SAFB bound genomic regions and their association with a predominantly repressive histone modification state, we further assessed our gene expression data to investigate whether genes associated with promoter-proximal or distal HOXA9/SAFB peaks showed differential directionality of expression. In both cases, the majority of genes were up-regulated upon perturbation of *HOXA9* and *SAFB* (Supplemental Figure 10), with myeloid differentiation-associated genes enriched in both groups (data not shown). Taken together, these data strongly indicate that HOXA9 together with SAFB imposes transcriptional repression of genes associated with myeloid differentiation through NuRD and HP1γ binding and activity.

### NuRD and HP1γ inactivation phenocopy HOXA9/SAFB at the functional and transcriptional level

To determine the functional relevance of NuRD complex and HP1γ co-occurrence at HOXA9/SAFB bound genomic regions, we targeted them genetically and via pharmacological inhibition of their obligate/recruited catalytic components. The NuRD complex contains obligate histone deacetylase activity and HP1γ interacts with the SUV39H1 protein that deposits H3K9me3, a mark read by heterochromatin (HP1) proteins. By treating the AML cell lines with a combination of a specific inhibitor of SUV39H1 (Chaetocin) and a pan-HDAC inhibitor (Panobinostat) we sought to further dissect mechanisms underlying HOXA9/SAFB mediated gene repression. Of note, both Chaetocin and Panobinostat have already been reported to inhibit the growth of AML cells alone or more potently in combination ^34–36^, however the gene expression programs and the chromatin-associated mechanisms that underlie these findings were not elucidated.

Remarkably, we could demonstrate that co-treatment with Panobinostat (here after referred as Pano) and Chaetocin, at non-cytotoxic concentrations in AML cell lines ^35,36^ could indeed phenocopy perturbation of HOXA9 or SAFB. In the HOXA-dependent MOLM13 and OCIAML3 AML cell lines the combination treatment caused a significant reduction in cell growth, along with the induction of apoptosis and differentiation, compared to the cells treated with single agents or vehicle-treated (Figures. 7A-D (MOLM13) and 7F-H (OCIAML3), Supplemental Figure 11A and 11C). As inhibition of HDAC and SUV39H1 activity is likely to have widespread transcriptional consequences, we specifically assessed the ability of the inhibitors to de-repress expression of nine candidate HOXA9/SAFB-repressed genes by RT-QPCR, and found similar de-repression of these genes as was observed upon HOXA9 or SAFB knockdown (Figure 7E and 7I, Supplemental Figure 11B and 11D). To further corroborate these findings, we also performed genetic perturbation of NuRD subunits (*MTA2* and *GATAD2A*) and *HP1*γ via CRISPR mediated knockout in MOLM13, and OCIAML3 cell lines. Genetic loss of these proteins also induced myeloid differentiation and apoptosis in both AML cell lines (Figure 7J, K and Supplemental Figure 12A). Finally, we examined the effect of genetic loss on expression of the same candidate genes by RT-QPCR, where we could demonstrate de-repression of the majority of genes following perturbation (Figure 7L, M and Supplemental Figure 12B and 12C).

**Figure 7.**
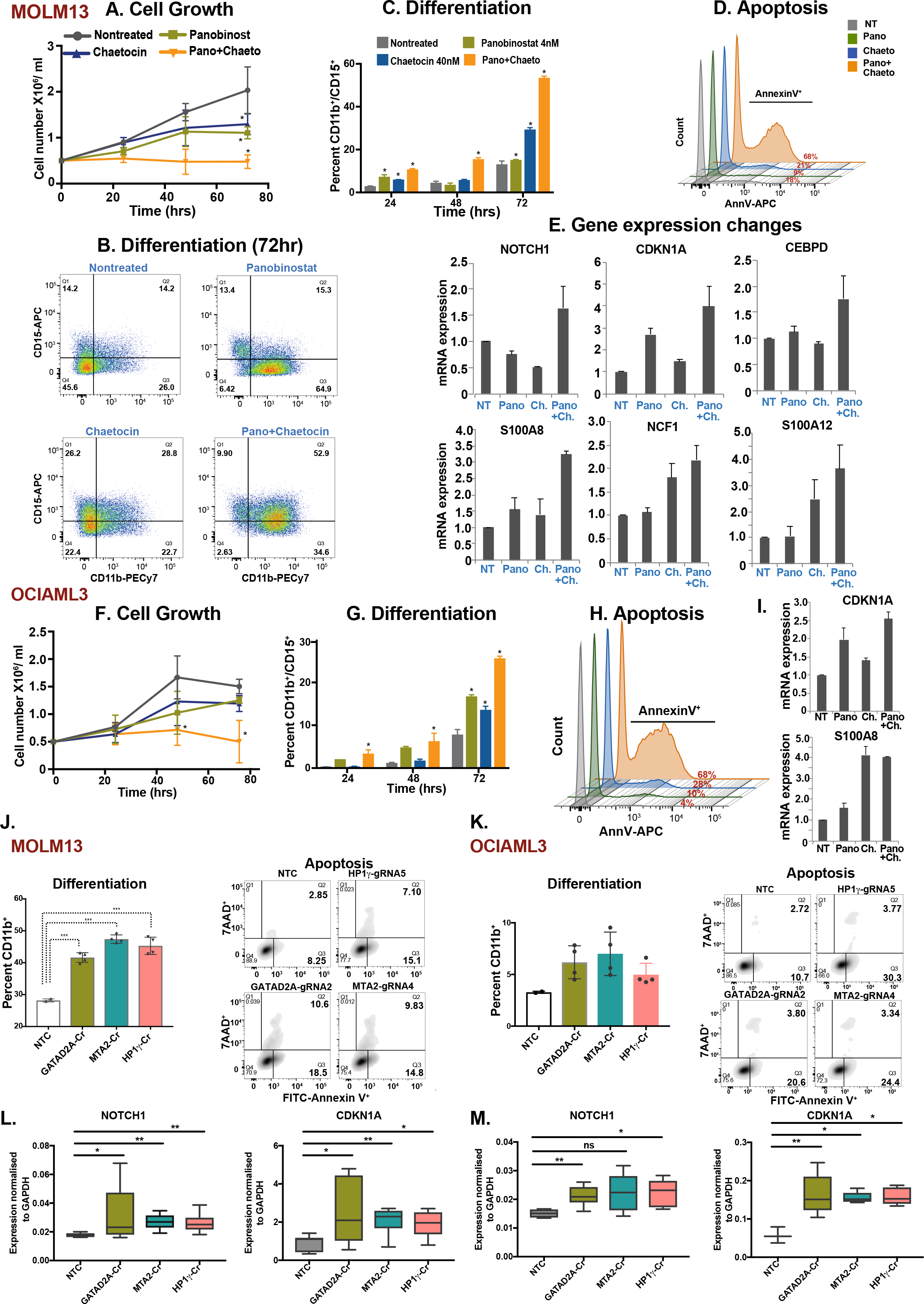
NuRD and HP1γ inactivation phenocopy HOXA9/SAFB at the functional and transcriptional level. A) Growth kinetics of MOLM13 cells treated with Pano (4nM), Chaetocin (40nM) alone or in combination. The data are shown as the average of biological replicates (n=3) ± SD. 2way ANOVA test, * p<0.01 comparing nontreated Vs treatments at 72hr. B) Scatter plot shows flow-cytometric analyses of CD11b-PECy7 and CD15-APC surface expression in MOLM13 cells treated with Pano (4nM), Chaetocin (40nM) alone or in combination. Representative plots of 3 independent biological replicates are shown. C) Bar graph shows the differentiation measured with CD11b-CD15 surface expression in MOLM13 cells after treatment with Pano (4nM), Chaetocin (40nM) alone or in combination over the time course. Data shown are an average of 3 biological replicates ± SD. 2-way ANOVA test, * p<0.001. D) Histogram shows flow-cytometric analyses of AnnexinV positive MOLM13 cells 72 hrs after treatment with Pano (4nM), Chaetocin (40nM) alone or in combination. Representative plots of 3 independent biological replicates are shown. E) RT-QPCR expression levels of selected target genes in MOLM13 cells treated with drugs alone or in combination for 48hrs. NT: nontreated, Pano:Panobinostat, Ch.: Chaetocin, Pano+Ch.: Panobinostat+Chaetocin. Data shown is representative of 3 independent biological replicates. F) Growth kinetics of OCIAML3 cells treated with Pano (4nM), Chaetocin (40nM) alone or in combination. 50,000 cells were seeded followed by daily counting. The data are shown as the average of 3 biological replicates ± SD. Statistical significance calculated using 2-way ANOVA test, * p<0.01. G) Bar graph shows differentiation measured by CD11b-CD15 surface expression in OCIAML3 cells after treatment with Pano (4nM), Chaetocin (40nM) alone or in combination for the time course. Data shown as an average of 3 biological replicates ± SD. Statistical significance calculated using 2-way ANOVA test, ** p<0.001. H) Histogram shows flow-cytometric analyses of AnnexinV positive OCIAML3 cells 72 hrs after treatment with Pano (4nM), Chaetocin (40nM) alone or in combination. Representative plots of 3 independent biological replicates are shown. I) RT-QPCR expression levels of selected target genes in OCIAML3 cells treated with drugs alone or in combination for 48hrs. NT: nontreated, Pano:Panobinostat, Ch.: Chaetocin, Pano+Ch.: Panobinostat+Chaetocin. The data shown here is representative of 3 independent biological replicates. J) Left panel shows a bar graph of the differentiation measured by CD11b surface expression in MOLM13 cells after CRISPR-kd of *GATAD2A* (p=0.0038), *MTA2* (p=0.0095) and HP1γ (p=0.0048). The data are shown as mean±SD, n=3. Statistical significance was calculated using the t-test. Right panel, shows density plots for apoptosis measured by AnnexinV and 7AAD positive cells after CRISPR-kd of *GATAD2A, MTA2* and HP1γ. Representative plots of 3 independent biological replicates are shown. K) OCIAML3, same as described in J. L) RT-QPCR analyses to measure expression of selected target genes in MOLM13 cells after CRISPR-kd of *GATAD2A, MTA2* and HP1γ. The data shown is summarised from 3 independent biological replicates. Statistical significance calculated using Mann-Whitney test * p<0.01. ** p<0.001. M) RT-QPCR in OCIAML3 cells, description as above in L.

Taken together these data demonstrate, for the first time, the importance of gene repression mediated by HOXA9 for the maintenance of the leukemia phenotype, identify the HOXA9/SAFB complex as central to this repression and demonstrate mechanistically that this repression occurs through the activity of the NuRD complex and HP1γ recruitment and activity.

## Discussion

Despite our understanding of the central role of HOXA9 in activating a conserved transcriptional program ^8,14^ that drives proliferation and survival of leukemic cells ^15,26^, very little is known of the specific mechanisms of action of HOXA9 in AML. Here we present novel observations informed by a protein interactome of cellular and chromatin-associated HOXA9 in AML that provides novel evidence for the leukemogenic activity of this protein. The matrix binding protein SAFB (Scaffold/Attachment factor B) is an interaction partner of HOXA9. Using *in vitro* and *in vivo* functional assays, and in conjunction with genomic assays performed before and after perturbation of the complex, we demonstrate that this functional interaction represses gene expression, with this repression a requirement for the maintenance of AML. Mechanistically, we demonstrate that this repression occurs through the recruitment of the NuRD remodelling complex and heterochromatin protein HP1γ to HOXA9-SAFB *cis*-regulatory sites to impose transcriptional repression on genes that are involved in myeloid differentiation and cell survival. Functionally, SAFB loss phenocopies that of HOXA9 in leukemic cells and loss of either protein triggers differentiation and apoptosis *in vitro* and *in vivo* in AML cells (Supplemental Figure-13).

The function of HOXA9 as an oncogene in AML has been almost solely attributed to its ability to activate gene programmes associated with leukemia ^15^. However, repressive functions have previously been associated with HOXA9 for genes including the p16^INK4a^/p19^ARF^ locus, although mechanistic detail has been lacking ^37,38^. We provide evidence that maintenance of the leukemic phenotype is dependent on the repressive effects of HOXA9 and that this repressive function requires the SAFB protein. SAFB was first described based on its ability to bind scaffold attachment region DNA elements and the nuclear matrix. This scaffold/matrix attachment region (S/MAR) protein (1) binds to both DNA and RNA ^39^; (2) stabilizes pericentromeric heterochromatin through interactions with major satellite RNA ^40^ (3) regulates DNA damage and (4) interacts with proteins and complexes involved in chromatin architecture and regulates transcriptional activation and repression through poorly understood mechanisms ^41–43^. Our data suggest that HOXA9 and SAFB, in turn interact with the NuRD and HP1γ co-repressor complexes to repress the induction of genes critical for normal myeloid differentiation, including *NOTCH, CDKN1A, CEBP*δ, *S100A8, S100A12*. Suppression of NOTCH signalling has been previously described as a requirement for AML, although the mechanism underlying this repression was previously unknown. Similar to NOTCH reactivation in *MLL* - rearranged leukemias ^44,45^, we could demonstrate that restoration of *NOTCH*, and other targets such as *CEBP*δ, via either exogenous over-expression or knock-down of *HOXA9* or *SAFB*, could inhibit the growth of leukemia cells and induce differentiation and cell death.

In HOXA9/MEIS1 transformed murine AML, Hoxa9 binding to intergenic regions facilitates the establishment of *de novo* enhancers, in conjunction with CEBPA and the MLL3/4 complex, leading to expression of leukemia-specific genes ^15^. However, here we have focussed on the HOXA9 binding sites that overlapped with SAFB in human leukemic cells. One third of these regions were in distal, rather than promoter elements and in the majority HOXA9/SAFB binding was associated with gene repression. S/MAR regions have been shown to flank enhancers, where they may provide boundary function and/or localise *cis*-regulatory elements to specific regions of the nucleus ^46,47^. Of interest, H9SB co-bound regions possessed sequence characteristics of S/MAR and modifications characteristic of both repressive (H3K9me2, H3K9me3, H3K27me3) and activated chromatin (H3K4me3 and increased accessibility ^48^. The genes linked to these regions are also known to be readily activated during myeloid differentiation. We speculate that when bound by HOXA9 and SAFB, NuRD and HP1γ are recruited to these elements to mediate repression, and that this cellular state is enforced in AML. However, upon loss of HOXA9 expression, as occurs during normal myeloid differentiation, we hypothesise that the elements are poised for subsequent rapid activation facilitating further maturation (Supplemental Figure 13).

Studies in *Safb*-deficient mice suggest that it may have a role in hematopoiesis. Homozygous deletion resulted in increased pre- and post-natal lethality, related to multiple developmental defects during embryogenesis, including reduced erythropoiesis ^49^. However, mice did survive to term, albeit with severe growth retardation. Our study identifies the HOXA9-SAFB axis as important for the maintenance of the differentiation block in AML. Due to the central role of HOXA9 in normal hematopoiesis and HSC function, we expect that it cannot be safely therapeutically targeted. However, targeting the HOXA9-SAFB recruited NURD and HP1γ catalytic activity or specifically targeting the HOXA9-SAFB interaction in AML may be possible. Further work will be necessary to determine the role of the HOXA9-SAFB interaction in normal haematopoiesis and to elucidate the stoichiometry and structural nature of this interaction to see if it is amenable to intervention.

A prominent role for liquid-liquid phase separation has recently been described in transcriptional regulation ^50^. Of note, two of the proteins in our putative repressive complex, SAFB and HP1 have recently been demonstrated to form biological condensates through Liquid-Liquid Phase separation (LLPS) ^51–53^. Moreover, the NuRD complex member MBD2 is predicted to induce phase separation due to the presence of a large RGG-rich stretch at its N-terminus ^54^ and the HOXA9 protein also has an internally disordered domain. SAFB and HP1 liquid condensates promote and stabilise the formation of heterochromatin and it is tempting to speculate that NuRD and HP1γ are not classically recruited to these loci by HOXA9/SAFB, but form co-condensed complexes within liquid condensates at cis-regulatory regions. Again, future work will be required to determine any relationship of the repressive complex to LLPS.

In summary, our study suggests that maintenance of a differentiation block is an active process that is critical for the leukemic state. We further demonstrate that this differentiation block is, at least in part, mandated by a novel repressive HOXA9-complex, thereafter detailing the molecular mechanisms that underlie the function of this HOXA9-SAFB complex (Supplemental Figure 13). Finally, our study suggests that either the HOXA9-SAFB complex or its recruited effector proteins may be potential therapeutic targets.

## Supplemental data

Reagents

Supplemental Table 1: HOXA9-Proteome

Supplemental Table 2-HOXA9/SAFB-common genome wide binding

Supplemental Table 3-RNAseq in CRISPRed cells

Supplemental Table 4-Genes Commonly-regulated by HOXA9/SAFB

Supplemental Table 5-NuRD-CBX3-H9SB-Overlapped regions

## Acknowledgements

This study was carried out in the laboratory of B.J.P.H. with funding from Cancer Research UK (C18680/A25508), the European Research Council (647685), MRC (MR-R009708-1), the Kay Kendall Leukaemia Fund (KKL1243), Worldwide Cancer Research (WCR14-1069) and the Cancer Research UK Cambridge Major Centre (C49940/A25117), This research was supported by the NIHR Cambridge Biomedical Research Centre (BRC-1215-20014), and was funded in part by the Wellcome Trust, who supported the Wellcome – MRC Cambridge Stem Cell Institute (203151/Z/16/Z) and Cambridge Institute for Medical Research (100140/Z/12/Z). The views expressed are those of the authors and not necessarily those of the NIHR or the Department of Health and Social Care. J.B. is funded by the Wellcome Trust 4-Year MRes+PhD Programme in Stem Cell Biology and Medicine (218481/Z/19/Z). D.S. was a postdoctoral fellow of the Mildred-Scheel Organization, German Cancer Aid (111875). A.D.W. was funded by a grant from Blood Cancer UK (13005). G.S.V. is a CRUK Senior Cancer Research Fellow (C22324/A23015). For the purpose of Open Access, the author has applied a CC BY public copyright licence to any Author Accepted Manuscript version arising from this submission. Authors acknowledge Dr. Pieter Van Vlierberghe from Ghent University for providing NOTCH-ICN-GFP retroviral construct.

## Authorship contributions

S.A.S and B.J.P.H Conceptualization and Methodology, B.J.P.H Funding Acquisition, Supervision, Resources, Project Administration, Writing and Reviewing, S.A.S Investigation, Validation, Formal analysis, Data curation, Project administration and Writing original draft, J.B. Investigation CRISPR knockdown studies, Validation, G.G. Investigation – invivo experiments, E.M.-Software, Formal analyses-bioinformatics data analyses, A.W., R.A., J.E.H., Investigation-Proteomic screen, S.A., A.S.B., F.S. Investigation-proteomic screen, Validation, G.V. Resources, D.S., H.Y., S.H., Resources and Writing – Review & Editing.

“The authors declare no potential conflicts of interest.”

---

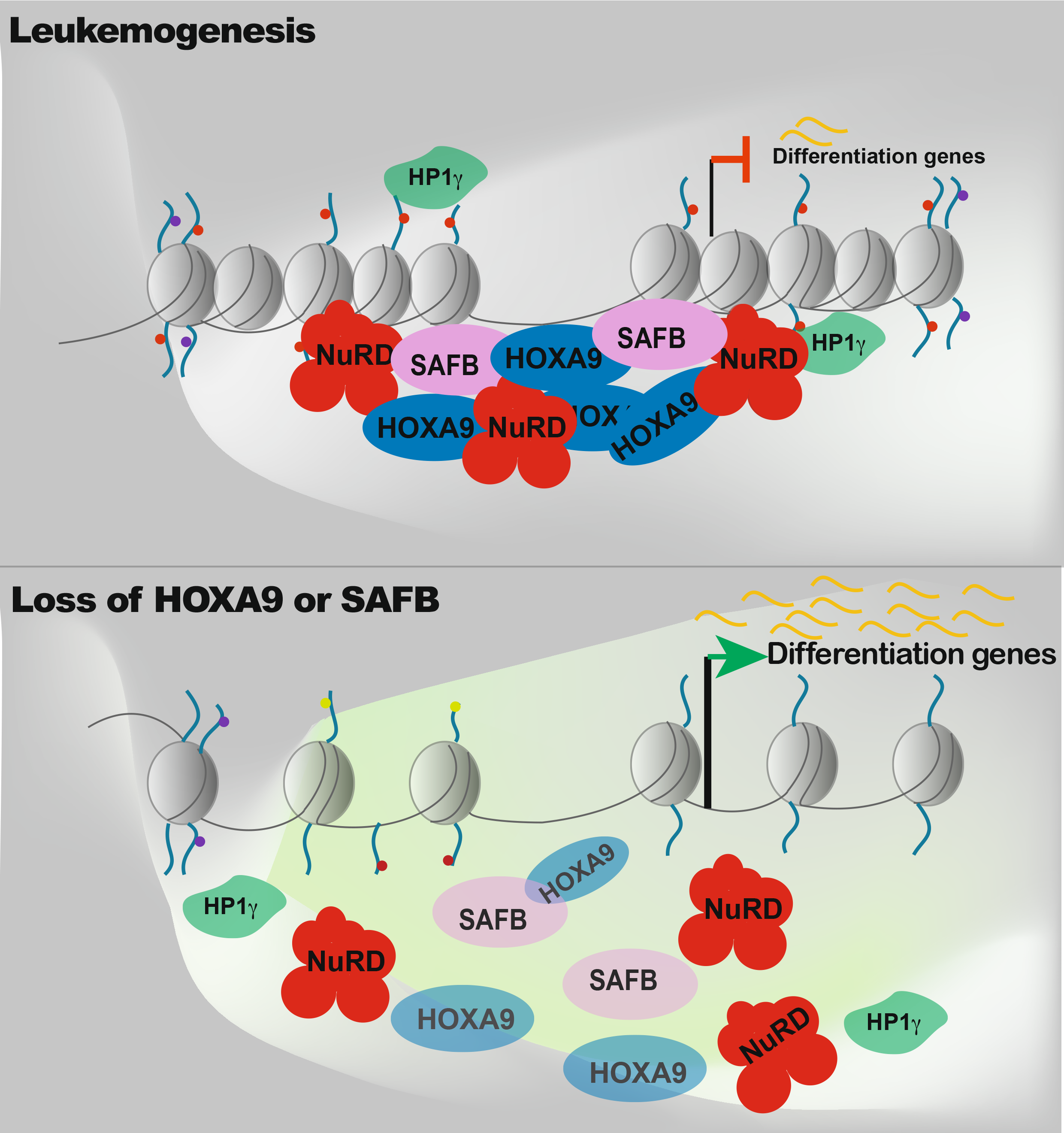

