## SupplementaryFiles for "HOXA9 forms a repressive complex with nuclear matrix-associated protein SAFB to maintain acute myeloid leukemia"

#### Supplementary Figure 1

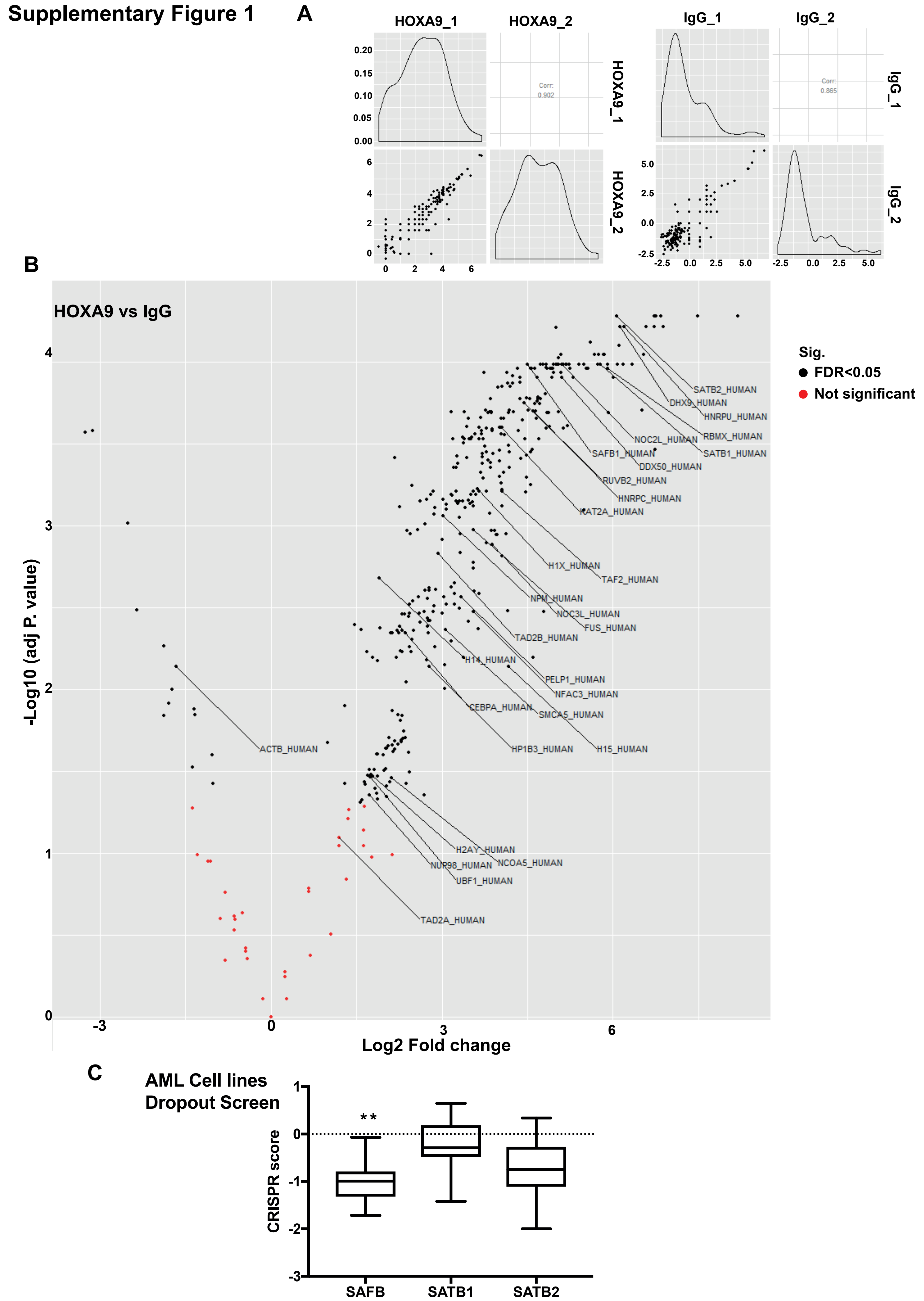

### Supplementary Figure 2

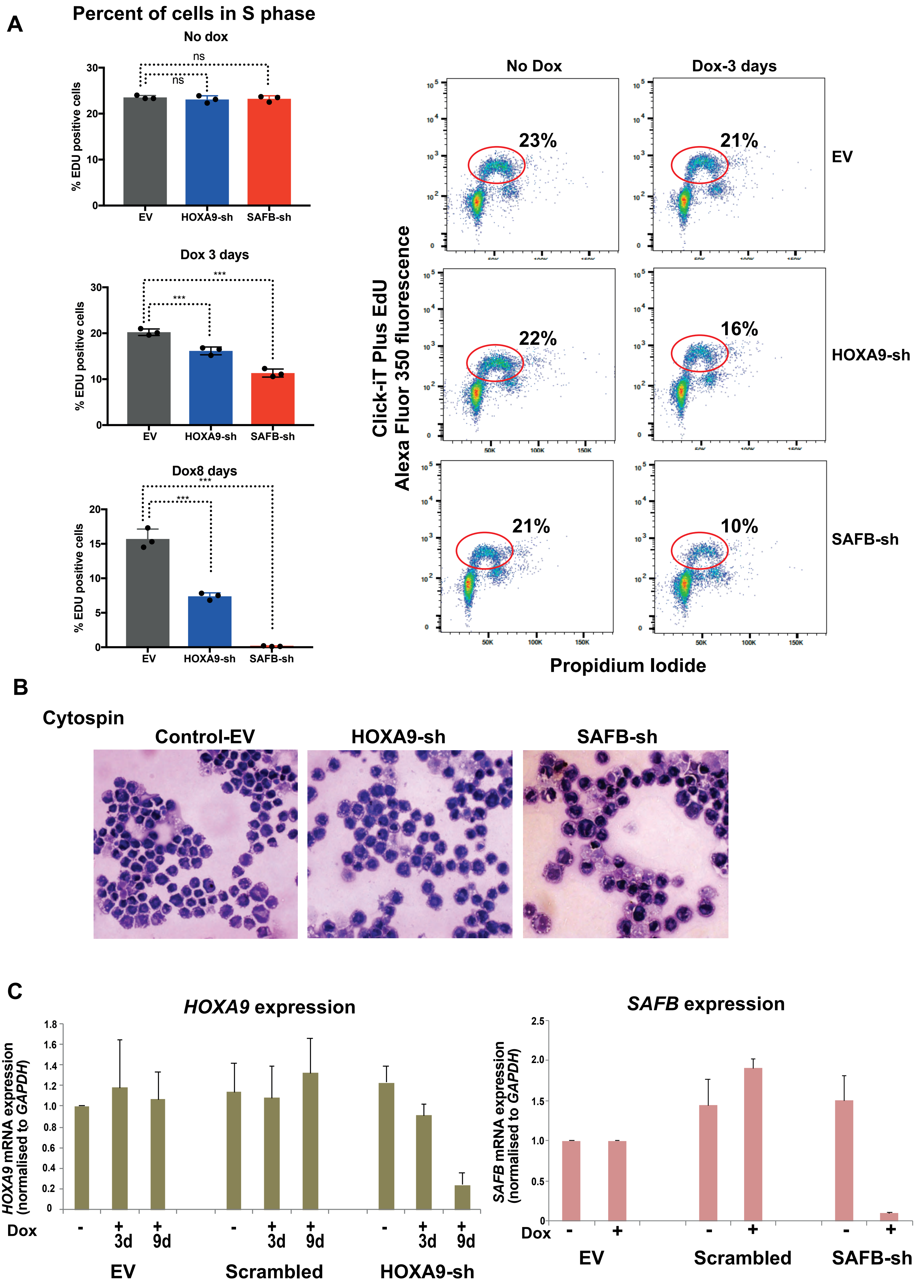

#### Supplementary Figure 3

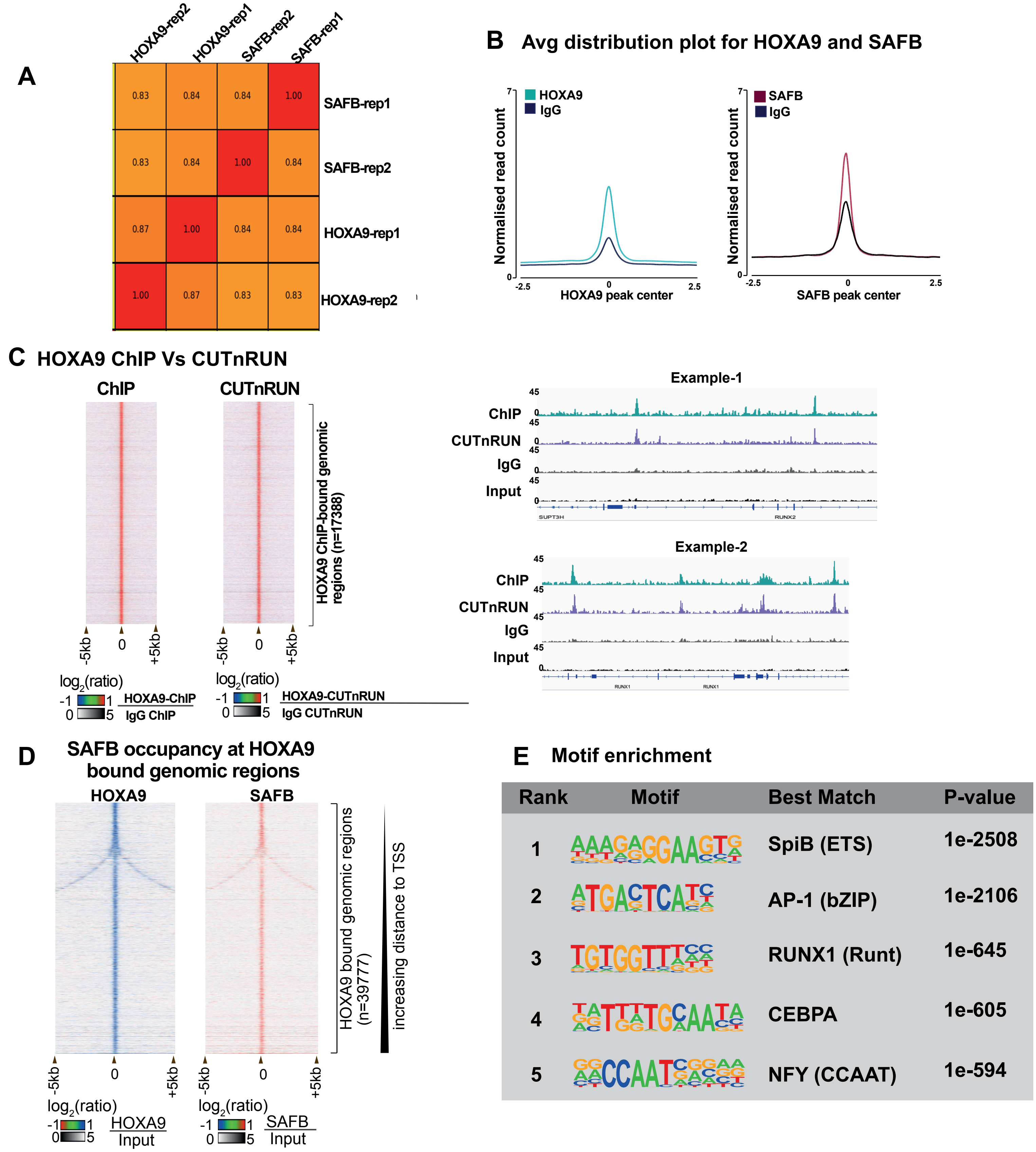

### Supplementary Figure 4

**A** S/MAR features on HOXA9-SAFB co-bound genomic regions

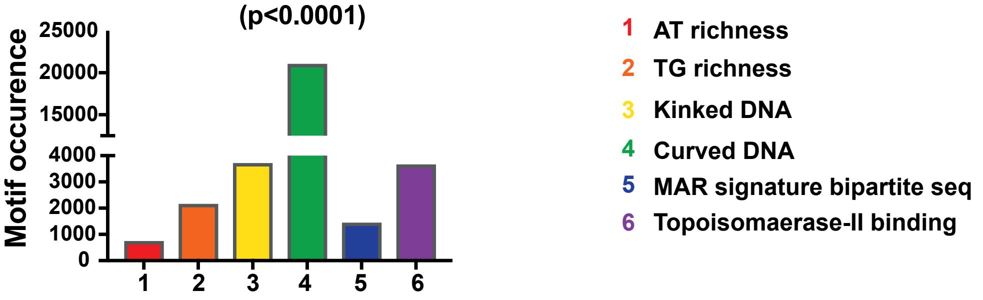

**B** HOXA9-SAFB occupancy at known classical S/MAR (shaded area)

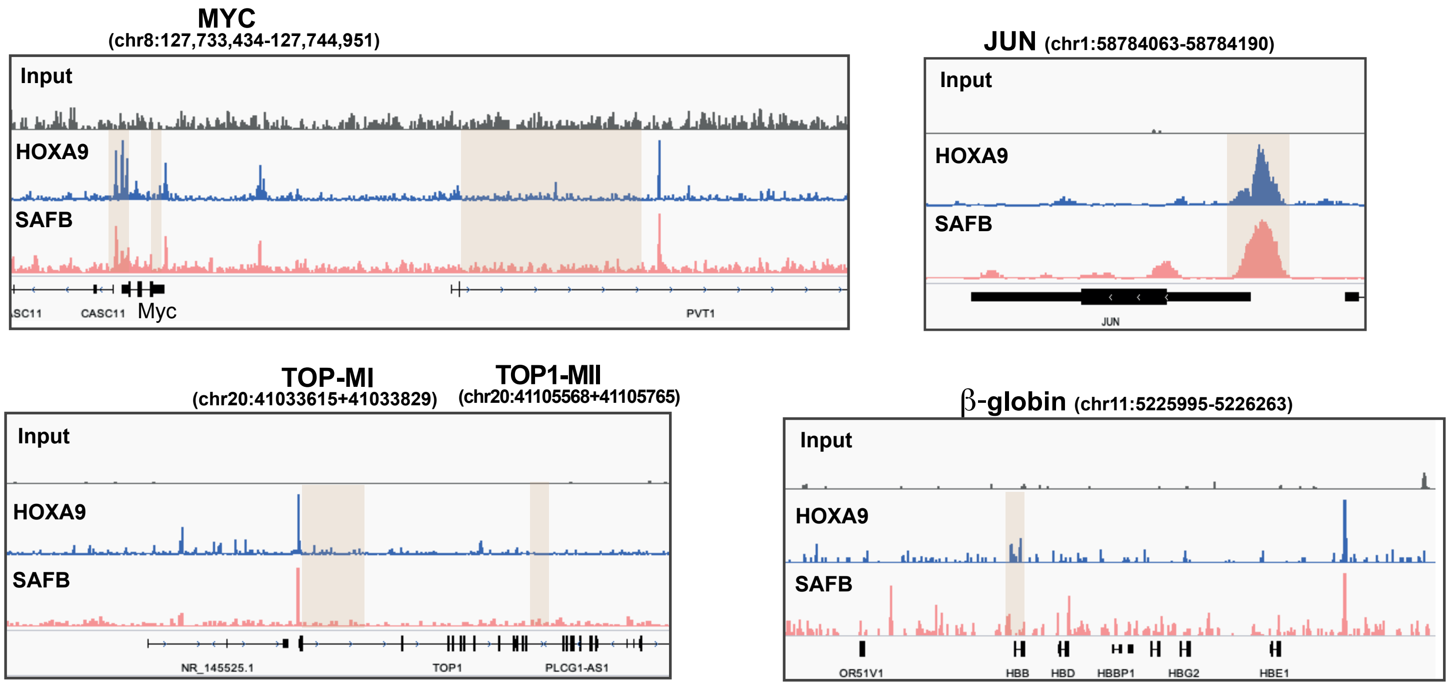

### Supplementary Figure 5

A

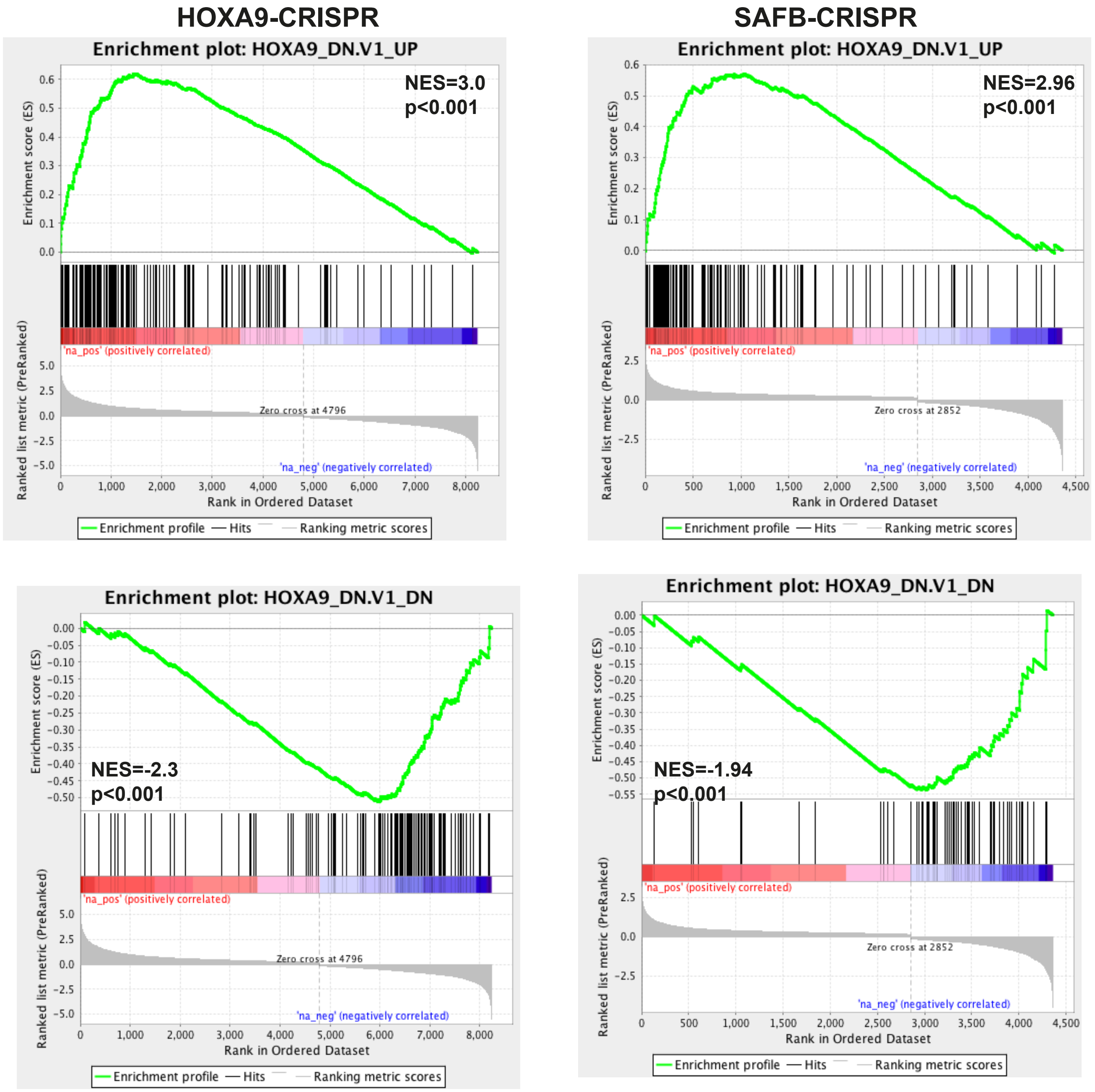

B

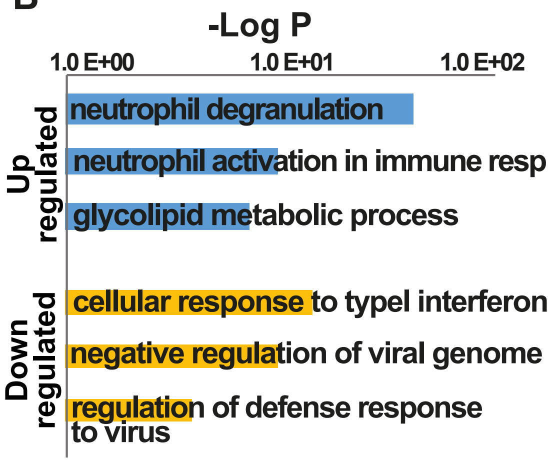

C

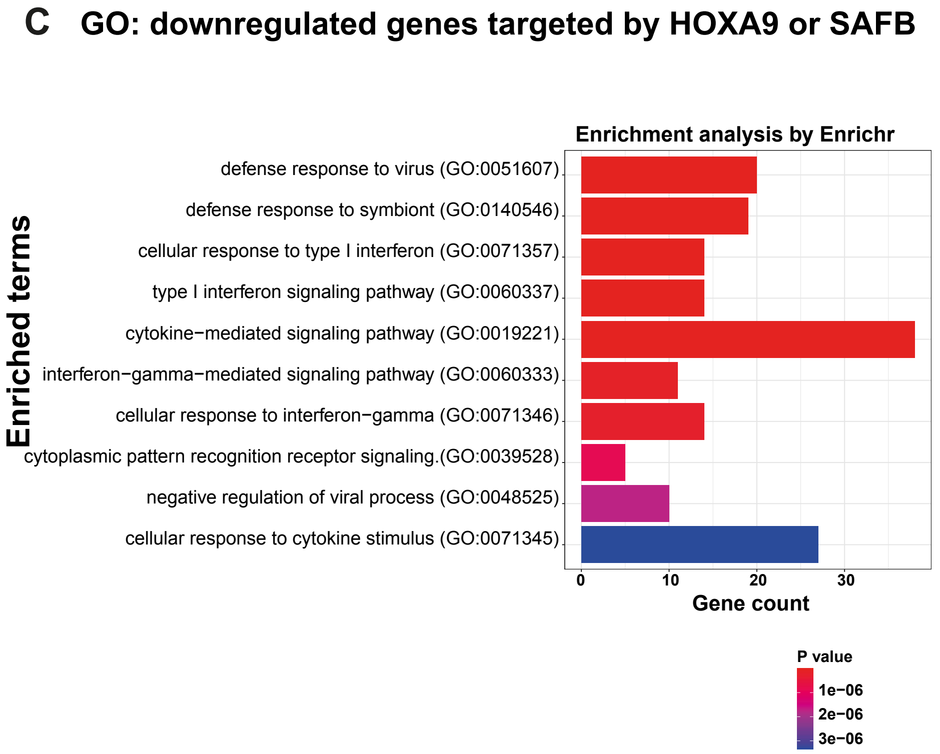

#### Supplementary Figure 6

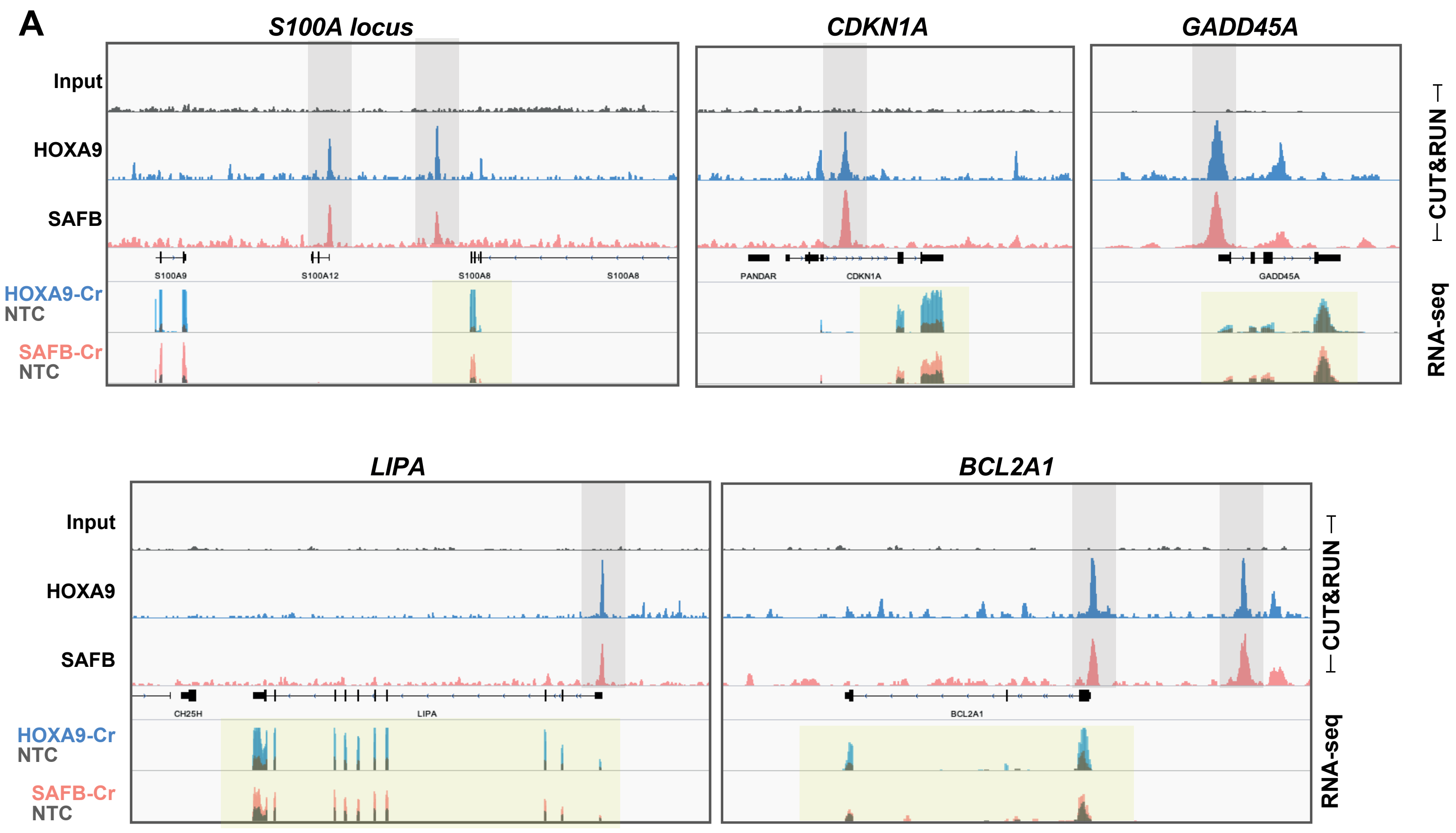

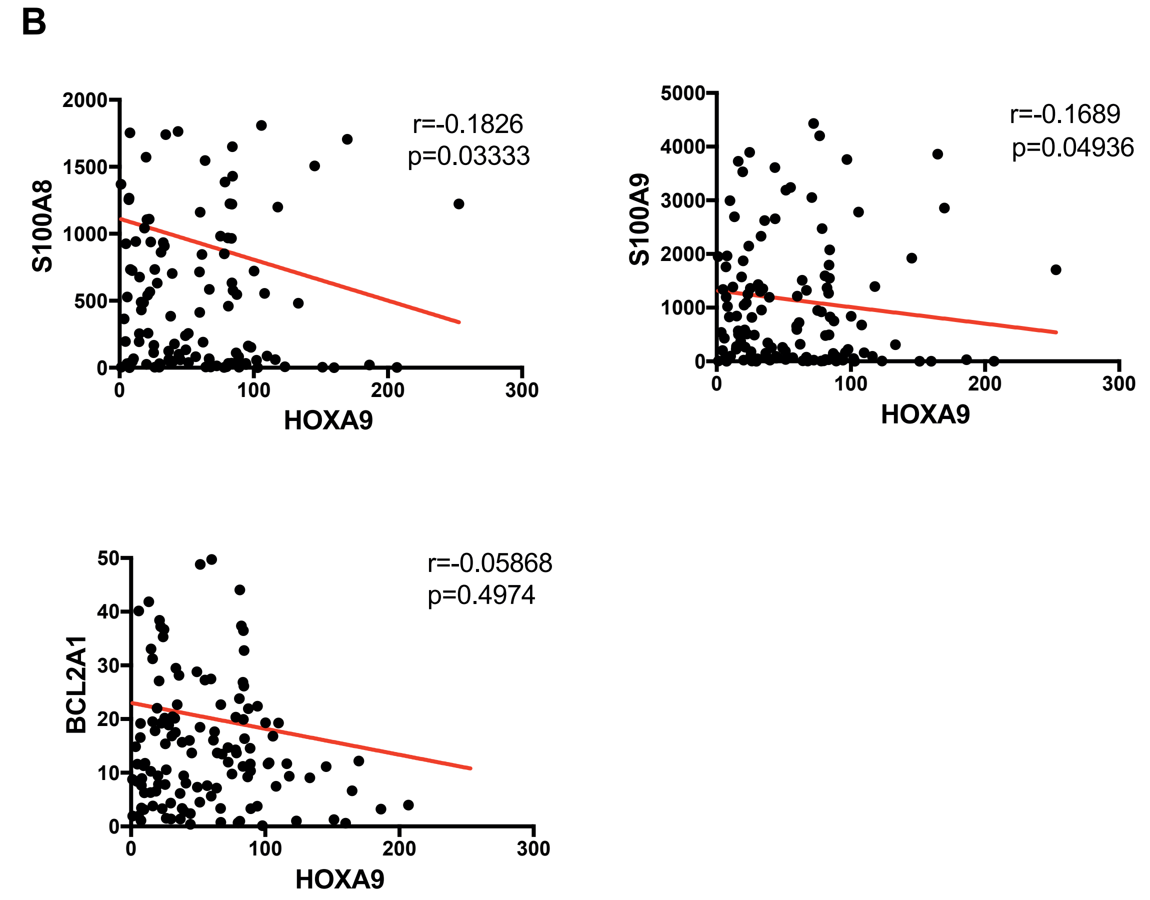

### Supplementary Figure 7

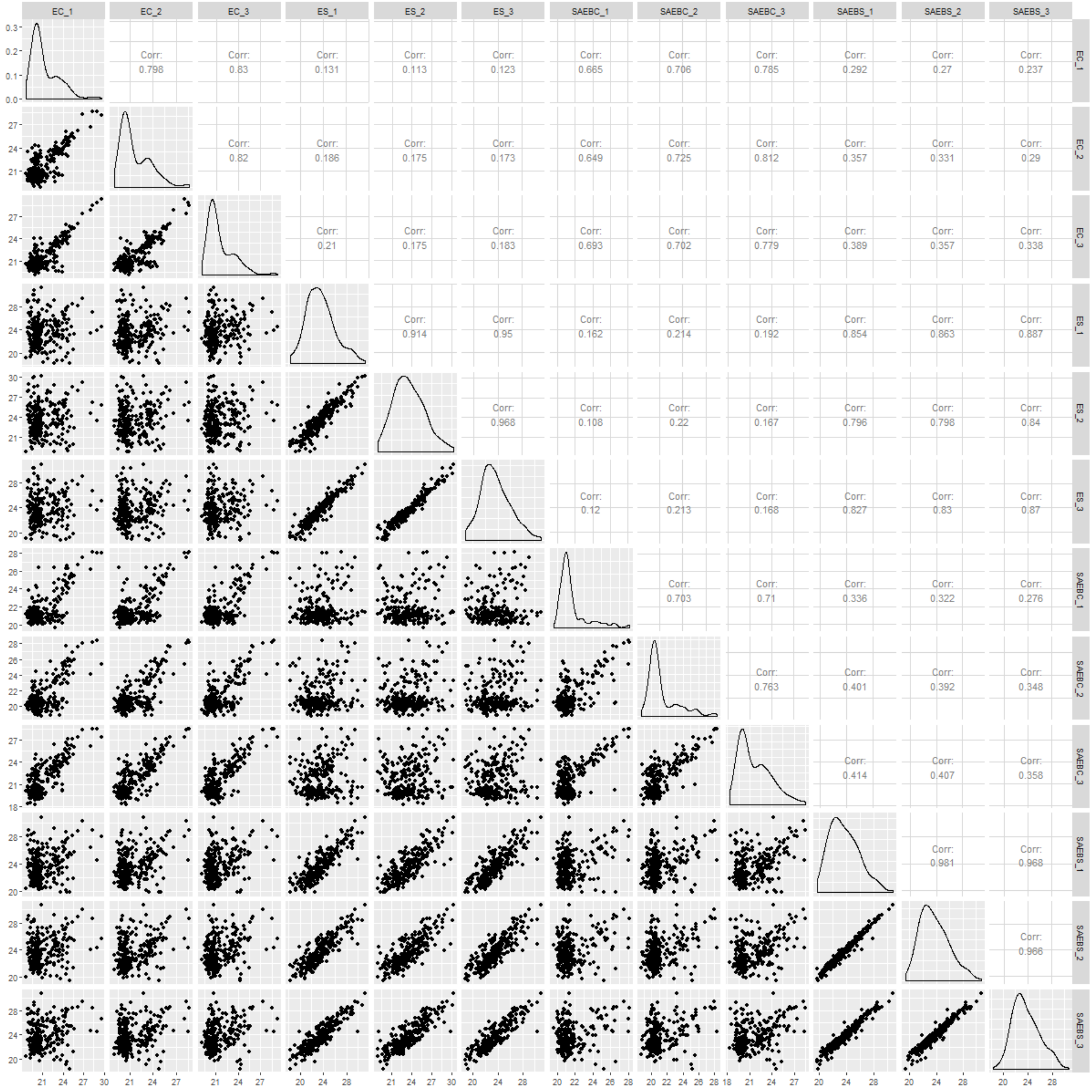

### Supplementary Figure 8

A

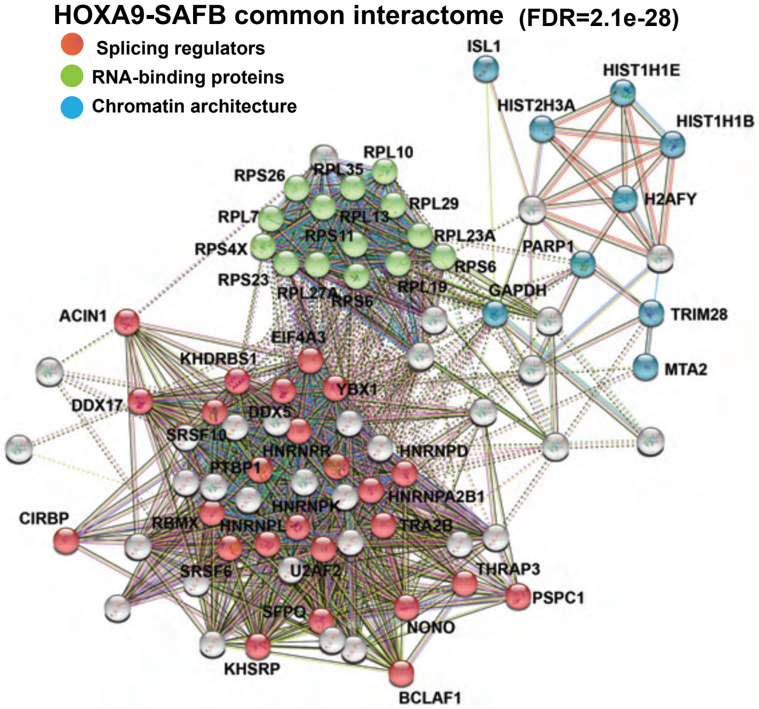

B

HP1γ (CBX3)-CUTnRUN replicates

GATAD2A-CUTnRUN replicates

MTA2-CUTnRUN replicates

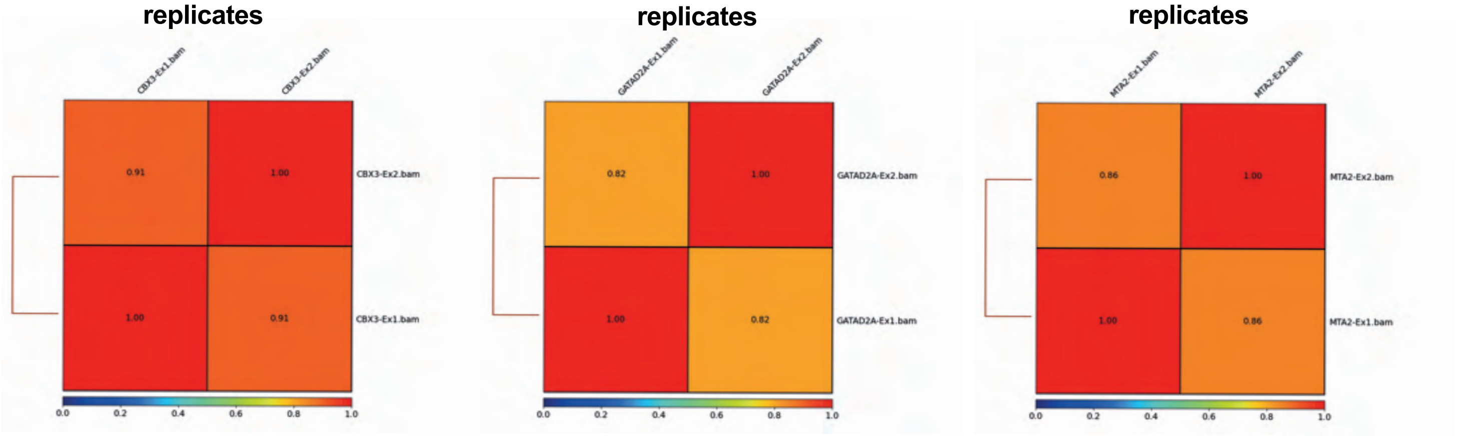

### Supplementary Figure 9

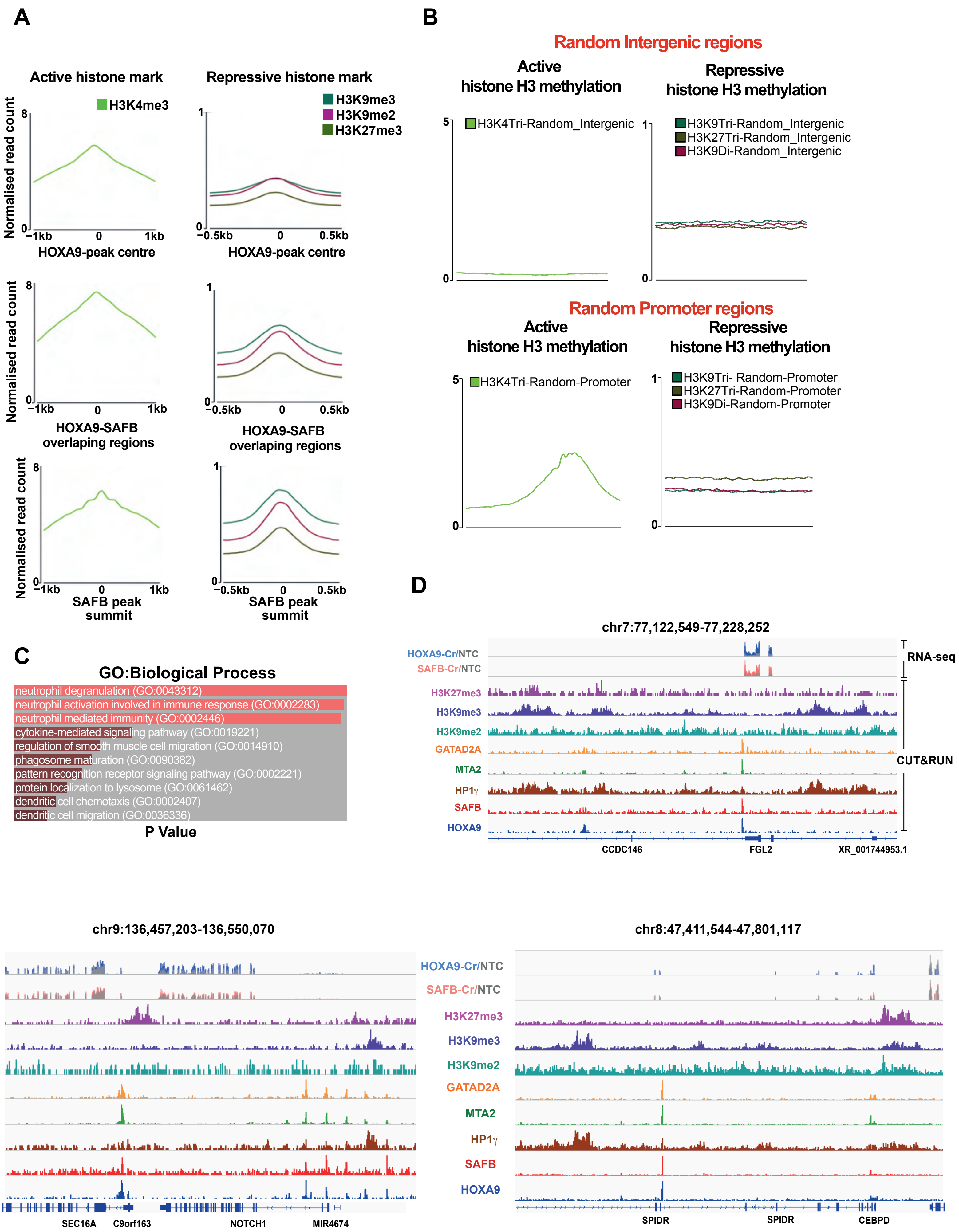

### Supplementary Figure 10

**A**

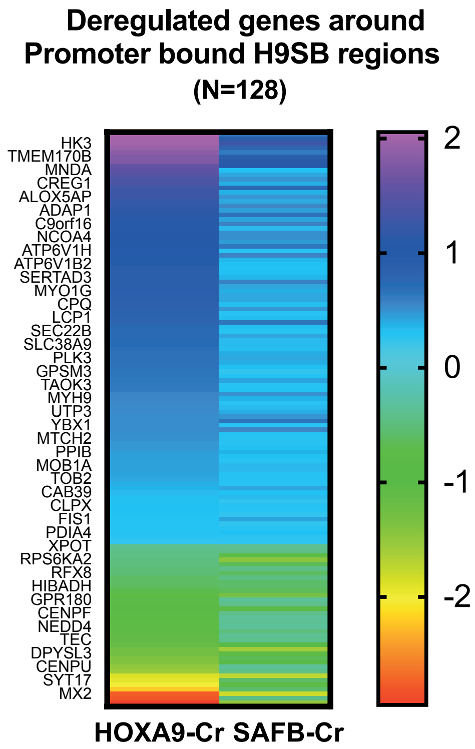

**B**

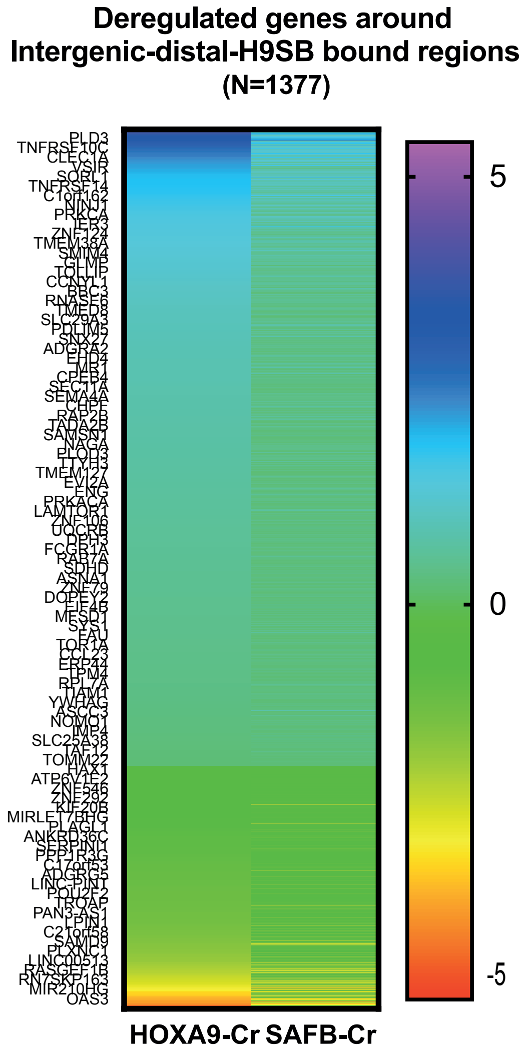

**Supplementary Figure 11**

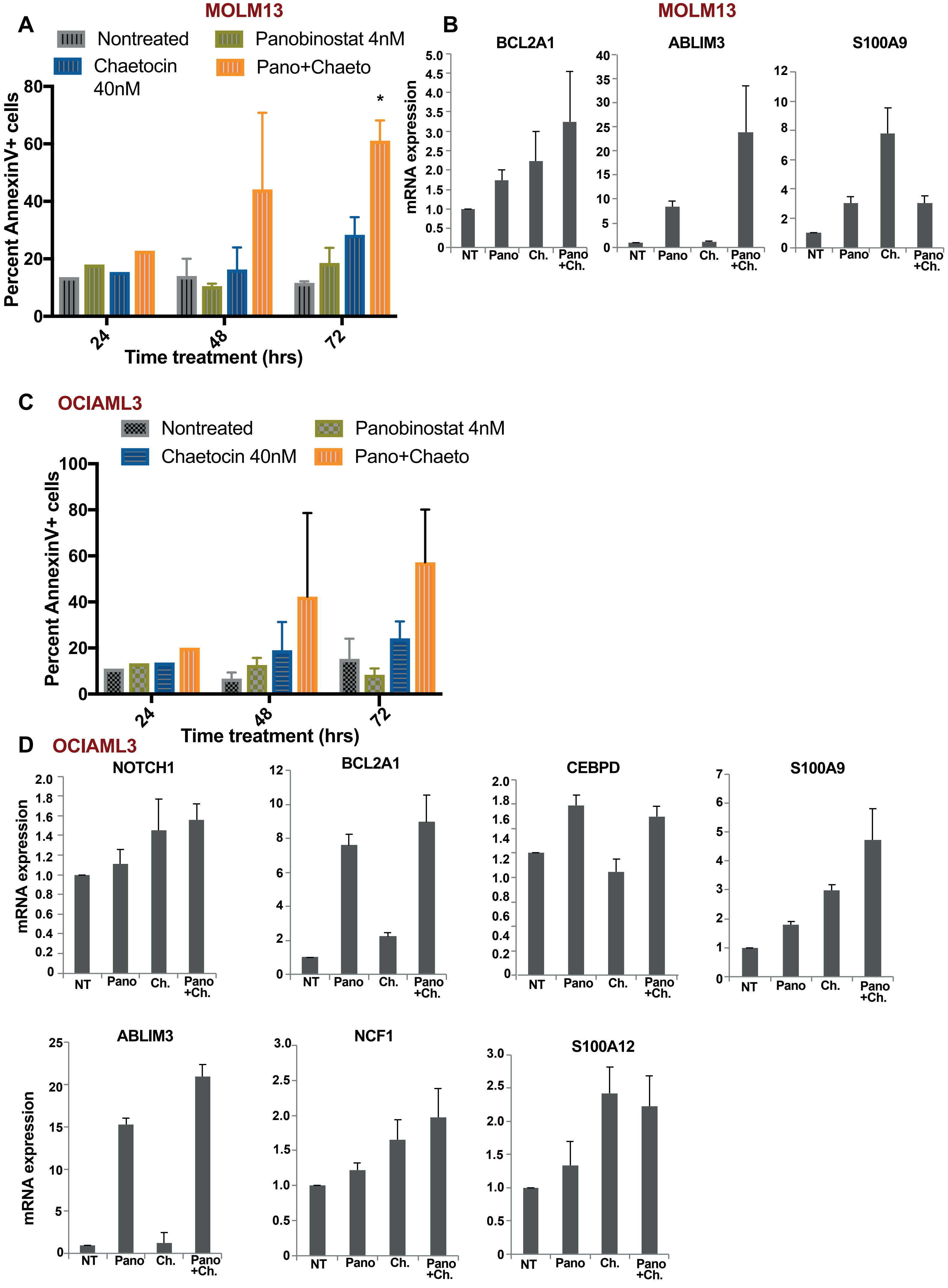

### Supplementary Figure 12

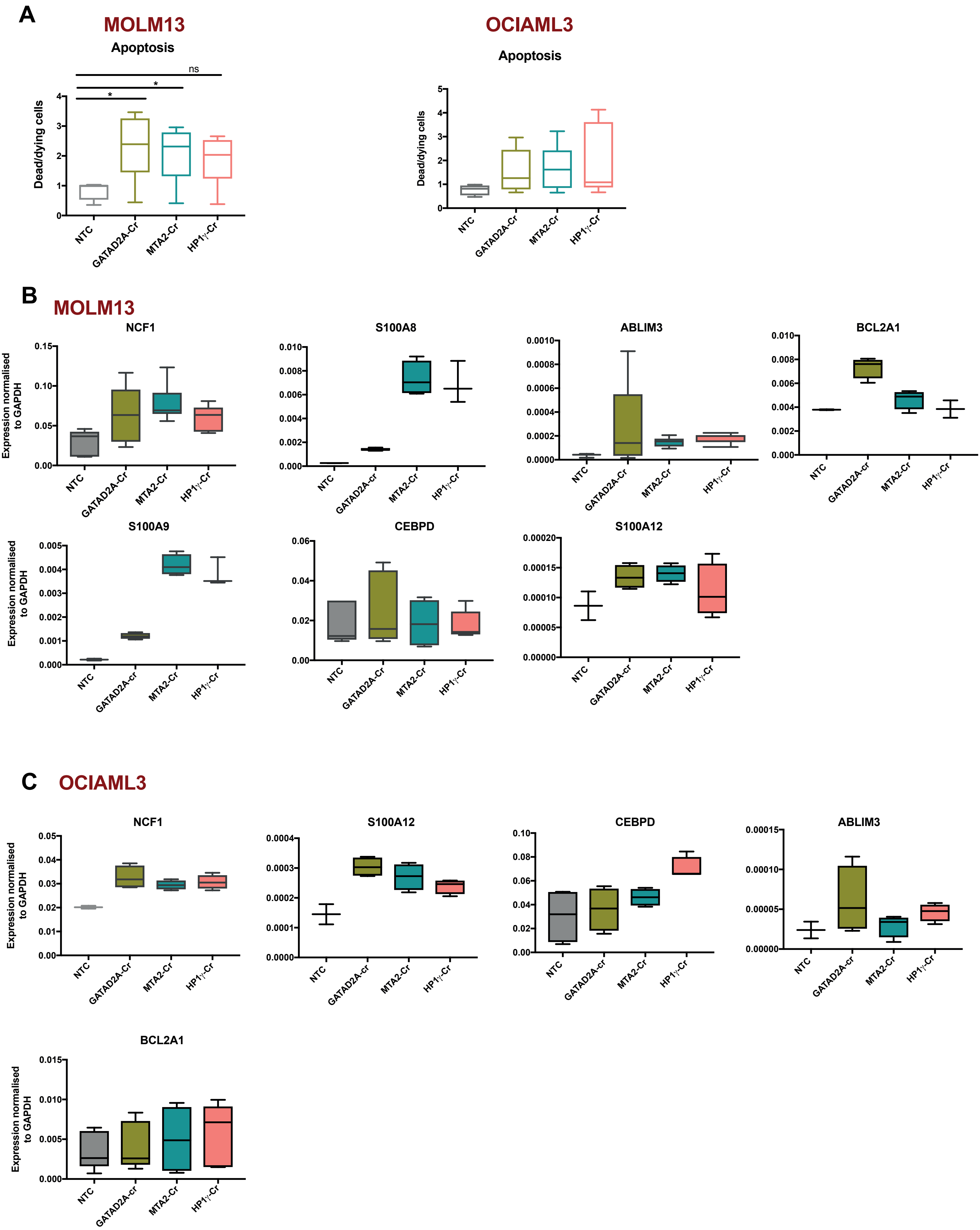

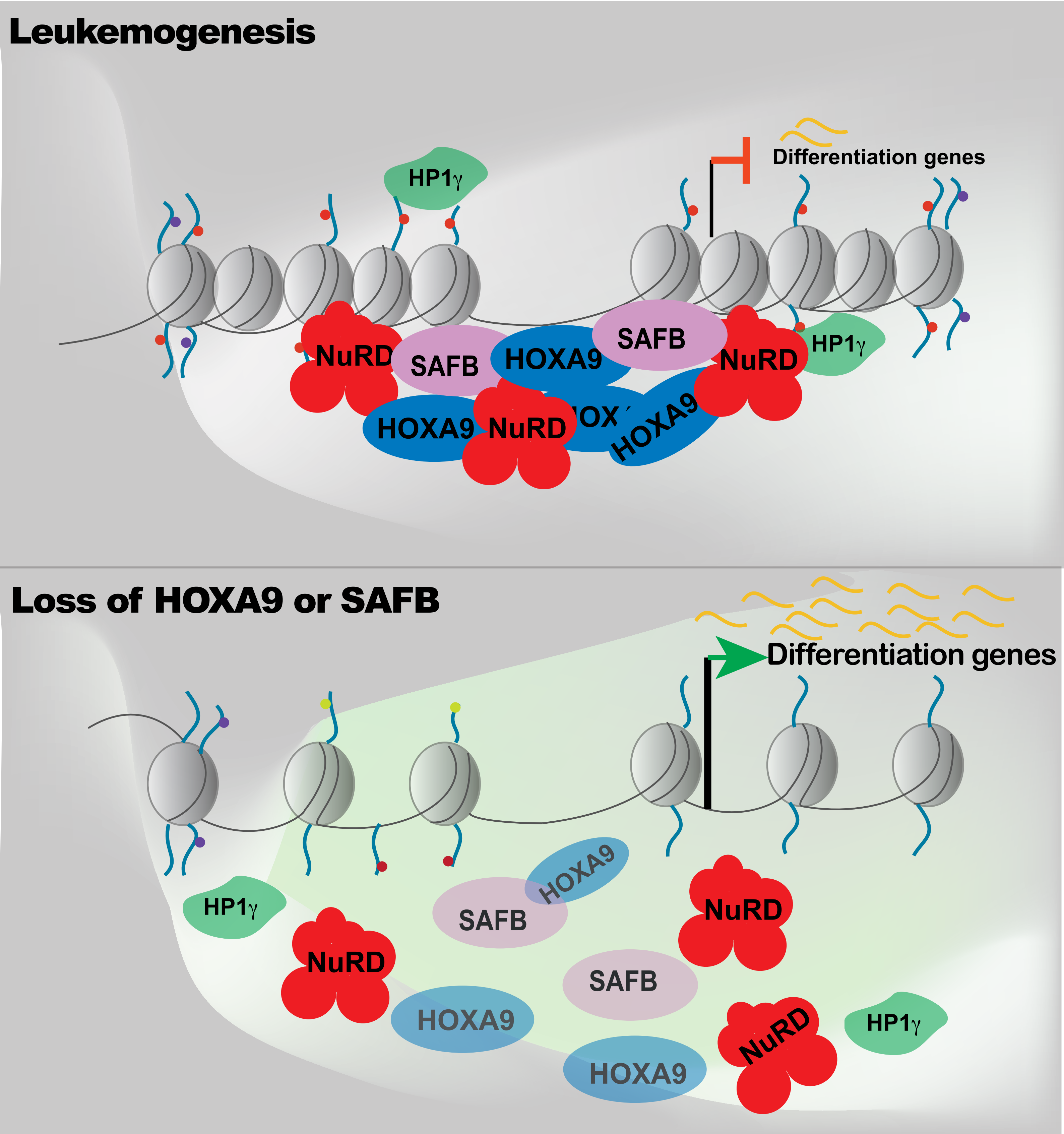
Leukemogenesis
HP1γ
Differentiation genes
NuRD
SAFB
HOXA9
SAFB
NuRD
HP1γ
HOXA9
NuRD
HOXA9
Loss of HOXA9 or SAFB
Differentiation genes
HOXA9
SAFB
NuRD
HP1γ
NuRD
SAFB
HOXA9
HOXA9
NuRD
HP1γ

**Supplementary Figure Legend**

**Supplementary Figure 1.**

A) The correlation between replicates (n=2) from mass spectrometric analyses of immunoprecipitated HOXA9 from MOLM13 cells is shown. The histograms show the distribution of protein abundances in each sample separately.
B) Volcano plot displaying label free quantitative MS results of HOXA9 pull down in MOLM13 cells. The plot shows log2 ratios of averaged peptide MS intensities between HOXA9-IP and control-IP (IgG) eluate samples (x axis) plotted against the negative log10 p values (y axis) calculated across the replicate data sets (one tailed Student's t test, n=2 replicates). Note maximum upper values were set for the x and y axis to accommodate all detected proteins in the plot. The full dataset of specific co-precipitating proteins is given in Table S1.
C) Box plot represents depletion of SAFB, SATB1, SATB2 by CRISPR in a dataset across 14 human acute myeloid leukemia (AML) cell lines (Wang et al., 2017).

**Supplementary Figure 2.**

A) The bar graph shows the percentage of S phase cells measured by Click-iT Plus EdU cell proliferation kit in shRNA expressing MOLM13 cells. A time course experiment was performed analysing cells at day3 and day8 of doxycycline treatment (1.5$\mu$g/ml).
B) Cytospins showing signs of myeloid differentiation in HOXA9 and SAFB shRNA cells.
C) RT-QPCR shows the expression of HOXA9 or SAFB after shRNA induction in MOLM13 cells.

**Supplementary Figure 3.**

A) Correlation matrix comparing peak overlap between the replicates of HOXA9 or SAFB enrichment from CUT&RUN sequencing in MOLM13 cells.

B) Average distribution plot shows the signal (intensity on Y-axis as normalised read count) for HOXA9 (blue) or SAFB (red) relative to IgG measured by CUT&RUN-sequencing in MOLM13 cells.
C) Heatmap comparison HOXA9 signal measured from ChIP or CUT&RUN using same antibody in MOLM13 cells. The Y-axis represents individual position of regions centred at HOXA9 peaks ±5kb around peak centre.
   Right panel: Two representative loci are shown for HOXA9 enrichment by ChIP and CUT&RUN in MOLM13 cells.
D) Heatmap shows the enrichment signal for SFAB at HOXA9 occupied genomic regions in MOLM13 cells by CUT&RUN.
E) Motif enrichment for HOXA9-SAFB co-occupied regions obtained from CUT&RUN by HOMER. Short 120bp fragments were used for motif search.

**Supplementary Figure 4.**

A) The bar graph shows presence of S/MAR features motif (Narwade et al., 2019) on the sequences obtained from HOXA9-SAFB co-bound genomic regions. Details are given in the method section.
B) Representative genomic loci taken from genome browser showing HOXA9-SAFB occupancy at known classical S/MAR (shaded area).

**Supplementary Figure 5.**

A) GSEA enrichment plot shows HOXA9 signature enriched in differential gene expression data from HOXA9 or SAFB perturbation in MOLM13 cells.
B) Gene ontology analyses for upregulated (upper panel, blue), downregulated (lower panel, yellow) genes. The FDR values are plotted as -logP on Y-axis.
C) Gene ontology analyses for downregulated genes in response to HOXA9 or SAFB perturbation in MOLM13 cells and possess proximal occupancy of HOXA9-SAFB (within 50kb to TSS). The gene count is plotted on X-axis. The colours on the bar represents the significance p-value. A bar represents the colour code is shown on the right side of the plot.

**Supplementary Figure 6.**

A) Representative loci from genome browser showing HOXA9 (blue) or SAFB (red) enrichment from CUT&RUN in MOLM13 cells. Grey shaded area shows colocalization of HOXA9 and SAFB. Lower part of the tracks shows the transcription of indicated genes measured from RNA-sequencing in MOLM13 cells with CRISPR knockdown of HOXA9 (blue) or SAFB (red) relative to nontargeting control (NTC, grey).
B) Correlation between *HOXA9* and *S100A8 or S100A9 or BCL2A1* mRNA expression in human AML primary samples (n=165, TCGA data set).

**Supplementary Figure 7.**

The correlation matrix showing high concordance between replicates (n=3) from mass spectrometric analyses of immunoprecipitated SAFB from MOLM13 cells. The histograms show the distribution of protein abundances in each sample separately. EC=Empty vector MOLM13 cells-IgG; ES= Empty vector MOLM13 cells-SAFB-IP; SAFB-C=SAFB-shRNA MOLM13 cells-IgG; SAFB-S=SAFB-shRNA MOLM13 cells-SAFB-IP.

**Supplementary Figure 8.**

A) The network displays the prediction of protein-protein interaction of HOXA9-SAFB interacting proteome. p value FDR =2.11e-28. K-mean clustering shows 3 clusters, Red (mainly contains splicing factor and RNA binding proteins), Green (Ribosomal proteins), Blue (chromatin bound gene repressors).
B) Correlation matrix comparing peak overlap between the technical replicates of CBX3, or GATAD2A, or MTA2 enrichment from CUT&RUN sequencing in MOLM13 cells.

**Supplementary Figure 9.**

A) Tracks of the histone modifications average signal (intensity on Y-axis as normalised read count) from CUT&RUN in MOLM13 cells centred at HOXA9-peaks (top panel) or HOXA9-SAFB-co-occupied regions (middle panel), SAFB-peaks (right panel).

B) Average signal for histone marks H3K4me3, H3K9me2, H3K9me3, H3K27me3, H3K9ac, H3K27ac at random genomic regions of similar size to HOXA9-SAFB bound genomic regions; intergenic (top panel), random promoters (lower panel).

C) Gene ontology analyses for genes upregulated (n=420) in response to HOXA9 or SAFB perturbation in MOLM13 cells and possess proximal occupancy of HOXA9-SAFB, NuRD and HP1$\gamma$ (within 50kb to TSS).

D) Genome browser track shows the co-localization of HOXA9/SAFB repressive complex, correlation with repressive histone modifications obtained from CUT&RUN sequencing in MOLM13 cells and the derepression of the gene upon HOXA9 or SAFB perturbation in RNA-seq data using Hg38 genome. FGL2, NOTCH1, and CEBPD locus.

**Supplementary Figure 10.**

A) The heatmap shows the expression Log2 fold change in (HOXA9-Cr Vs NT) or (SAFB-Cr vs NT) MOLM13 cells for genes present in the vicinity of HOXA9-SAFB cobound regions; promoter bound (top panel),

B) distal/intergenic region (lower panel). A bar showing the relationship between colouring and expression value is shown at the right side of the plot.

**Supplementary Figure 11.**

A) The bar graph shows percentage of apoptotic MOLM13 cells (AnnexinV positive) after treatment with Panobinostat, or Chaetocin alone or in combination in a time course experiment. The data are shown as average of biological replicates (n=3) ± SD. Statistical significance was calculated using 2way ANOVA test, * p<0.01.

B) RT-QPCR to measure expression of selected target genes in MOLM13 cells treated with drugs alone or in combination for 48hrs. NT: nontreated, Pano:Panobinostat, Ch.: Chaetocin, Pano+Ch.: Panobinostat+Chaetocin. The data shown here is representative of 3 independent biological replicates.
C) The bar graph shows percentage of apoptotic OCIAML3 cells (AnnexinV positive) after treatment with Panobinostat, or Chaetocin alone or in combination in a time course experiment. The data are shown as average of biological replicates (n=3) ± SD.
D) RT-QPCR to measure expression of selected target genes in OCIAML3 cells treated with drugs alone or in combination for 48hrs. NT: nontreated, Pano:Panobinostat, Ch.: Chaetocin, Pano+Ch.: Panobinostat+Chaetocin. The data shown here is representative of 3 independent biological replicates.

**Supplementary Figure 12.**

A) The bar graph shows percentage of apoptosis (AnnexinV positive) in MOLM13 cells (left panel), OCIAML3 (right panel) after CRISPR knockdown of GATAD2A, MTA2, CBX3. The data are shown as average of biological replicates (n=3) ± SD. Statistical significance was calculated using 2way ANOVA test, * p<0.01, **p<0.001.
B) RT-QPCR to measure expression of selected target genes in MOLM13 cells after CRISPR knockdown of GATAD2A, MTA2, CBX3. The data are shown as average of biological replicates (n=3) ± SD, a median line represents the frequency of expression.
C) RT-QPCR to measure expression of selected target genes in OCIAML3 cells after CRISPR knockdown of GATAD2A, MTA2, CBX3. The data are shown as average of biological replicates (n=3) ± SD, a median line represents the frequency of expression.

Narwade, N., Patel, S., Alam, A., Chattopadhyay, S., Mittal, S., and Kulkarni, A. (2019). Mapping of scaffold/matrix attachment regions in human genome: a data mining exercise. Nucleic Acids Res *47*, 7247-7261. 10.1093/nar/gkz562.
Wang, T., Yu, H., Hughes, N.W., Liu, B., Kendirli, A., Klein, K., Chen, W.W., Lander, E.S., and Sabatini, D.M. (2017). Gene Essentiality Profiling Reveals Gene Networks and Synthetic Lethal Interactions with Oncogenic Ras. Cell *168*, 890-903 e815. 10.1016/j.cell.2017.01.013.

#### Methods

##### Cell Lines

MOLM13, MV411, HL60 cells were cultured in RPMI1640 medium containing 10%FBS, 1%L-Glutamine and 1%PenStrep. OCIAML3 cells were cultured in MEMa medium containing 20%FBS, 1%L-Glutamine and 1%PenStrep. Cas9 expressing OCIAML3 and HL60 were received from Prof. George S. Vassiliou, University of Cambridge. MOLM13 and MV411 cells with stable Cas9 expression were generated by using lentiCas9-Blast vector (Addgene#52962). Single cell clones were obtained by culturing cells in methylcellulose media followed by picking single colonies and established in liquid culture. Cas9 activity and expression was confirmed in multiple clones independently.

293T cells were cultured in DMEM medium containing 10% (v/v) FBS, 1% (v/v) L-Glutamine and 1% (v/v) PenStrep. All cell lines were routinely tested negative for mycoplasma contamination.

For treatment with inhibitors Panobinostat-LBH589 (Cat # S1030, Selleckchem) and Chaetocin (Cat # HY-N2019, MedChemExpress), cells were seeded in 6 well plate at density of $5X10^5$/ml in 3 ml media. Inhibitors were diluted to working stock of 10mM with DMSO and were used at final concentration of 4nM Panobinostat and 40nM Chaetocin. Cells were harvested for assays at specific time points.

##### Cloning of sgRNA into lentiviral vector

The chosen sgRNAs sequences were designed with overhang sequence for BbsI restriction sites. Annealing of the sense 1 µl (100 pmol/µl) and antisense 1 µl (100 pmol/µl) gRNA oligonucleotides for each gRNA was performed in 10x NEB buffer 2: 2 µl in a final volume of 20 µl, with denaturation at 95°C for 5 min in a heating block and slowly allowed to anneal oligonucleotides until mixture is at room temperature (which takes approximately 1-2 h). The annealed gRNA oligonucleotides can be store at -20 °C for later use or used for ligation, all in one tube by assembling reaction in a PCR tube:

i. 100 ng pkLV2 gRNA expression plasmid

ii. 1 µl of annealed gRNA oligonucleotide

iii. 1 µl BbsI restriction enzyme

iv. 1 µl T4 DNA ligase.

v. 2 µl 10x T4 ligase buffer (to a final concentration of 1x).

vi. Nuclease-free water up to 20 µl total reaction volume.

The reaction was incubated in a thermal cycler for 10 cycles for [5 min at 37 °C, 10 min at 22 °C], hold for 30 min at 37 °C followed by 15 min at 75 °C, then hold 4°C. Competent *E. coli* bacterial was transformed using 2 µl of the ligation product and colonies were sequenced to confirm the clones using pKLV2 sequencing primer AGATAATTAGAATTAATTTGACTG.

**Cloning cDNAs into lentiviral vector**

NOTCH-ICN-GFP retroviral construct was a kind gift from Dr. Pieter Van Vlierberghe from Ghent University.

Human CEBPD cDNA was ordered from GenScript (NM_005195.4) ORF clone (Catalog no. OHu17203D). Primers were designed to PCR out the CEBPD cDNA with overhangs to clone with EcoRI and BamHI sites into pLVX-TetOnePuro vector (Clontech). Clones were confirmed by sanger sequencing.

**Cloning shRNAs into lentiviral vector (pLKO-TetOn)**

shRNA against human HOXA9 and SAFB were cloned into pLKO "all-in-one" system for the inducible shRNA expression as described earlier [17,18].

***In vivo* transplantation of human leukemia cells**

MOLM13 cells were transduced with lentiviruses expressing the puromycin resistance gene and a doxycycline-inducible shRNA against *HOXA9* or *SAFB* or a control (scrambled) shRNA. Cells were selected by 2µg/ml puromycin. Knockdown was confirmed by RT-QPCR and western blotting. The selected shRNA stable cells were further engineered to express luciferase for bioluminescence studies. GFP positive cells were sorted and maintained as stable sh-luciferase-MOLM13 cells. For *in vivo* studies, 100,000 cells were transplanted via tail vein injection into sub-lethally irradiated (2Gy) 6-8 weeks old female NSG (NOD.Cg-Prkdcscid Il2rgtm1Wjl/SzJ) mice. Disease dissemination was confirmed and subsequently tracked by IVIS bioluminescence imaging (PerkinElmer). In brief, D-luciferin (Cat#122799, Perkin Elmer) was administered by intraperitoneal (IP) injection (10ul/gr of body weight of a 15mg/ml (DPBS) solution) followed by inhalation anaesthesia (isoflurane) and IVIS bioluminescence imaging (seven-and-a-half minutes post luciferin injection for all animals/imaging sessions). Tumour burden was quantified using Living Image Software (version 4.7.2, PerkinElmer). To induce shRNA expression, mice were provided with a doxycycline containing feed (Cat# A112D72003, Ssniff Spezialdiäten GmbH). All mice were housed in a pathogen-free animal facility and were allowed unrestricted access to food and water. All experiments were conducted under a UK Home Office project (under the Animals (Scientific Procedures) Act 1986, Amendment Regulations (2012)) and following ethical review by the University of Cambridge Animal Welfare and Ethical Review Body.

**Co-Immunoprecipitation**

Co-immunoprecipitation was performed in MOLM13 cells using Nuclear Complex Co-IP Kit (Active Motif, Cat#54001) following manufacturer's instructions. The immunoprecipitated samples were processed for mass spectrometry as described below or were run on 4-15% gradient SDS-PAGE gels (Biorad) and transferred to PVDF membrane and followed for western blotting in western blotting section using HRP-conjugated light chain specific secondary antibody (Cat# 211-002-171).

**Rapid immunoprecipitation mass spectrometry of endogenous proteins (RIME)**

RIME was performed as described earlier (Mohammed et al., 2016). Briefly, shRNA containing MOLM13 cells (SAFB-sh, HOXA9-sh, Scrambled) were treated with doxycycline (1.5ug/ml) for 5 days. The knockdown and differentiation phenotype were confirmed before fixing the cells with formaldehyde. 60million cells were fixed with 1% (v/v) formaldehyde and followed the procedure as described in the paper (Mohammed *et al.*, 2016). Antibodies used for RIME were HOXA9 (Atlas Antibodies, Cat# HPA061982), SAFB (Millipore, Cat# 05-588) or Rabbit IgG control (Proteintech, Cat# 30000-0-AP). Three independent replicates were analysed for SAFB or IgG antibodies whereas two replicates were performed for HOXA9 due to limited availability of HOXA9 antibody. We only considered SAFB-associated or HOXA9-associated proteins that were present in all independent replicates for corresponding pull-downs and excluded any protein that occurred in any one of IgG control RIME. The MaxLFQ values were analysed to compare different samples, that includes standard statistical testing combined with quantification accuracy for each of the quantified proteins across different samples [19].

##### Mass spectrometry (MS) based proteomics analyses

Digestion was performed in 2steps; step1) for 30 minutes at 27°C and 800rpm/min Urea buffer plus 5ug/ml Trypsin, followed by 2 washes with Urea buffer plus 1mM DTT. Step 2) overnight at room temperature with the pooled supernatants from step1 and washes. Next day, peptides were alkylated for 30 minutes with freshly prepared, light protected Iodoacetamide (IAA, SigmaDAldrich) solution and acidified with Trifluoroacetic, acid, (TFA, Sigma5Aldrich). Desalting was next performed using in house prepared staged tips with Solution A (0.1% TFA in H2O) and solution B (50% Acetonitrile (ACN, SigmaDAldrich), 0.1% TFA in H2O), while elution was performed prior to loading into a DionexRSLC3000UPLC on line to a Thermo Orbitrap Q Exactive mass spectrometer

##### Data Processing

Raw files were analysed with MaxQuant (version 1.6.6.0) with integrated Andromeda used for searching MS spectra. The following parameters were used: specific trypsin digestion; up to 2 missed cleavages allowed; Precursor ion tolerance: 20 and 4.5 ppm for first and main searches, respectively; Human database (UniProt reference proteome downloaded 18 Dec 2018 containing 21066 proteins) with additional inbuilt MaxQuant contaminant database (containing 246 common contaminants); oxidation (M) and N terminal acetylation as variable modifications; carbamidomethylation (C) as a fixed modification; label-free quantification enabled (MaxLFQ algorithm), LFQ min. ratio count = 2; Match between runs enabled. Proteins were filtered for FDR (< 1%), reverse sequences and contaminants (both excluded), and three valid values required. Missing values were imputed from a normal distribution (0.3 width, 1.8 SD downshift. Post-processing was carried out using R with bespoke scripts incorporating the "limma" statistical package.

Protein interaction network analyses were performed using STRING online tool (https://string-db.org). The prediction is based on direct (physical) or indirect (functional) associations. STRING version 11.0 was used for this analysis. Number of nodes 79, average local clustering classification 0.716, PPi enriched p value <1.0e-

16, number of edges 1095, 48 proteins fall into negative regulation of gene expression category (GO0010629).

**Western blotting**

Cells were counted and washed twice with cold PBS prior to collection. One million cells were resuspended in 30ul 2XDTT buffer [1.24g DTT, 4ml 20% SDS, 4ml Glycerol, 0.4ml 10% Bromophenol Blue, for total volume of 40ml with water], vortexed and boiled for 5 min. Whole cell lysates were run on 4-15% gradient SDS-PAGE gels (Biorad) and transferred to PVDF-FL membrane. Membranes were blocked in 5%BSA in TBS-tween for 30 min at room temperature (RT), and incubated with primary antibody overnight at 4°C, followed by washes in TBS-Tween and incubation with secondary antibody-fluorescent tag (LI-COR) for 30min. Blots were scanned in LI-COR instrument.

**Lentiviral production**

Lentiviral particles were produced in 293T cell lines with co-transfecting packaging plasmids (pMDG2 and psPAX2) with pkLV2 (gRNA expression vector) or pLKO-Tet-on (shRNA vector) using TransIT transfection reagent. The supernatant containing viral particles were collected at 36 hrs post transfection in IMDM medium containing 30%FBS and filtered through 0.35um filters and stored at -80°C.

**Generation and validation of CRISPR-KO cell lines**

CRISPR knockout was performed using 4-5 distinct sgRNA via lentiviral delivery of guide RNA into cells stably expressing spCas9 (Addgene #52962). The lentiviral plasmid used for cloning sgRNA was pKLV2-U6gRNA5(BbsI)-PGKpuro2AmCherry-W or BFP (Addgene #67977). Briefly, cells were freshly seeded at 1million/ml density in 6 well plate in 3ml medium. One ml of lentivirus was added onto cells (1:4 dilution), with 8-10ug/ml polybrene. Cells were spun at 3000rpm for 1.5hr at 32°C. Following spinoculation, cells were incubated overnight at 37degree. Next day, medium was changed with fresh media. And cells were harvested for subsequent experiment on following day.

**Colony forming unit assay (CFU)**

CRISPR-ko or shRNA expressing MOLM13 cells were seeded (1000cells per ml) in methylcellulose (MethoCult H4434, Stem Cell Technologies). CFU were scored and photographs were taken using STEMVision instrument (Stem Cell Technologies).

**Cumulative growth curve**

Cells were seeded at 50,000 cells/ml density. For shRNA containing cells, expression was induced with doxycycline (1.5µg/ml). Cells were counted daily or alternate days depending on the experiment. Cells were maintained at initial density every 2 days. Fold accumulation was calculated by comparing to the starting cell number, followed by multiplying the fold change each consecutive day. The data shown as average of biological replicates (n=3) ± SD. Statistical significance was calculated using prism7 software, 2way ANOVA test.

**Competition assay**

Competition assay was performed between Flow sorted GFP positive NOTCH-ICN or MSCV-control vector expressing MOLM13 cells. Equal number of GFP positive cells were seeded at day0 and GFP percentage was measured upto 5 days by flowcytometry. Data shown as average of biological replicates (n=3) ± SD.

**Cumulative cell growth**

Cell growth of MOLM13 cells with CEBPδ overexpression with or without doxycycline was performed on stable cell lines (M13-CEBPδ or M13-Empty vector). Cells were seeded at density of 50,000 cells per ml density and counted alternate days. Cell density was maintained at initial concentration after each count and fresh dox was added to the culture. Fold accumulation was calculated by comparing to the starting cell number, followed by multiplying the fold change each consecutive day. The data shown as average of biological replicates (n=3) ± SD. Statistical significance was calculated using prism7 software, 2way ANOVA test. Error bars represent ± SD.

**Flow cytometry (FACS) analysis**

To monitor the differentiation status, cells were stained with the following antibodies PECy7-αHu-CD11b (Cat#301322, Biolegend), APC-αHu-CD15 (Cat#17-0158-42, eBioscience), FITC-αHu-CD14 (Cat#301804, Biolegend), [1:100 in FACS buffer: PBS with 2%FBS]. To measure apoptosis, cells were washed once with Annexin V binding buffer, and incubated with anti-FITC-AnnexinV (Cat# 640945, Biolegend) and 7-AAD (Cat# 420404, Biolegend) in the AnnexinV binding buffer for 15 min based on manufacturer's instructions. Cells were analysed on a BD-Fortessa instrument. Data were analysed using FlowJo software version 10.6.1.

**Cell Cycle analyses**

Cell cycle analyses was performed using Click-iT Plus EdU cell proliferation kit (Invitrogen, Cat# C10645) following manufacturer's instructions. Cells were analysed on a BD-Fortessa instrument. Data were analysed using FlowJo software version 10.6.1.

**RNA isolation and qPCR**

Total RNA was extracted from cells using RNeasy Plus mini (QIAGEN) kit following manufacturer's instructions. For RNA-seq, libraries were prepared using Illumina kit (NEBNext Ultra II Directional RNA Library Prep Kit for Illumina, Cat# E7760S) with RiboZero depletion following manufacturer's instructions. Libraries were sequenced on Hiseq platform with PE150 reads length.

For RT-QPCR, 1ug of RNA was reverse transcribed into cDNA using Taqman Reverse Transcription Reagents kit (applied biosystem, Cat# N8080234). QPCR was performed using an Mx3000P qPCR System (Agilent) detection system using primers (sequences listed in primers table) together with SYBR green master mix (Brilliant III Ultra Fats SYBR green QPCR master mix, Agilent, Cat# 600882).

**CUTnRUN**

CUTnRUN was performed using CUTANA CUT&RUN kit (Epicypher) following manufacturer's instructions. 100,000 MOLM13 cells were used for the assay for each antibody. Antibodies are listed in the table. For Input, 100,000 cells were used and treated with MNase similar to antibody containing sample and processed as other samples. Libraries were prepared using 2ng of DNA using NEBNext UltraII DNA library preparation kit (Cat# E7645S, Illumina) and Dual index primer-Multiplex Oligos (Cat# E7600S, Illumina). The samples were amplified for 12 cycles using KAPA HiFI 2X master mix (Cat# 07958935001, Roche). After amplification library was size selected using Ampure beads. The library size and concentration were analyzed using

Tapestation reagents-D5000 (Cat# 5067-5588, 5067-5589, Agilent) in 4200 TapeStation system (Agilent) before pooling the samples. The samples were sequenced at Novaseq SP50bpPE.

Details about bioinformatic analyses are provided in supplemental methods.

All sequencing data are deposited to GEO

#### gRNA Sequences

| gRNA-Name | Sequence |
|---|---|
| SAFB_gRNA1_S | CCGCAGATCGATCACTCGC |
| SAFB_gRNA1_AS | GCGAGTGATCGATCTGCGG |
| SAFB_gRNA2_S | AACGGAATGTGGACTCGAG |
| SAFB_gRNA2_AS | CTCGAGTCCACATTCCGTT |
| SAFB_gRNA3_S | GAAATTGAAATTACCTCCG |
| SAFB_gRNA3_AS | CGGAGGTAATTTCAATTTC |
| SAFB_gRNA4_S | ACGGGCTGGAGGAAAACTC |
| SAFB_gRNA4_AS | GAGTTTTCCTCCAGCCCGT |
| SAFB_gRNA5_S | CGGGCATTTGTCACAACCT |
| SAFB_gRNA5_AS | AGGTTGTGACAAATGCCCG |
| HOXA9 gRNA1-S | GAGCGTTGGCCGCTATGCGC |
| HOXA9 gRNA1-AS | GCGCATAGCGGCCAACGCTC |
| HOXA9 gRNA4-S | TTAATGCCATAAGGCCGGC |
| HOXA9 gRNA4-AS | GCCGGCCTTATGGCATTAA |
| HOXA9 gRNA5-S | AGCGCATGTACCTGCCGTC |
| HOXA9 gRNA5-AS | GACGGCAGGTACATGCGCT |
| HOXA9 gRNA6-S | GCGCCTGGGGGTGCACGTA |
| HOXA9 gRNA6-AS | TACGTGCACCCCCAGGCGC |
| HOXA9 gRNA7-S | TCGTGGAACCCAGTGCACG |
| HOXA9 gRNA7-AS | CGTGCACTGGGTTCCACGA |
| HOXA9 gRNA8-S | GCTATGCGCCGGGGACCCT |
| HOXA9 gRNA8-AS | AGGGTCCCCGGCGCATAGC |
| Nontargeting 1-S | ATTTTCGTACCCTGGGACGC |
| Nontargeting1-AS | GCGTCCCAGGGTACGAAAAT |

| | |
|---|---|
| MTA2-gRNA1-S | ATCCCAGATCGCCTAGTAG |
| MTA2-gRNA1-AS | CTACTAGGCGATCTGGGAT |
| MTA2-gRNA2-S | GTCCTGTACCGTATGTGGG |
| MTA2-gRNA2-AS | CCCACATACGGTACAGGAC |
| MTA2-gRNA4-S | TGGGGTACCAGGGTCGACA |
| MTA2-gRNA4-AS | TGTCGACCCTGGTACCCCA |
| MTA2-gRNA5-S | AACGGCTACGACCTGGCTA |
| MTA2-gRNA5-AS | TAGCCAGGTCGTAGCCGTT |
| GATAD2A-gRNA2-S | AACACGTGGCTCACCATGA |
| GATAD2A-gRNA2-AS | TCATGGTGAGCCACGTGTT |
| GATAD2A-gRNA3-S | GAGTGCCCCGAACAAGCGG |
| GATAD2A-gRNA3-AS | CCGCTTGTTCGGGGCACTC |
| GATAD2A-gRNA4-S | TCATGCCCCCACTCGTCAG |
| GATAD2A-gRNA4-AS | CTGACGAGTGGGGGCATGA |
| GATAD2A-gRNA5-S | AACATTGGCGACGCGGATG |
| GATAD2A-gRNA5-AS | CATCCGCGTCGCCAATGTT |
| CBX3-gRNA1-S | GAGCCTGAAGAATTTGTCG |
| CBX3-gRNA1-AS | CGACAAATTCTTCAGGCTC |
| CBX3-gRNA2-S | TAGATCGACGTGTAGTGAA |
| CBX3-gRNA2-AS | TTCACTACACGTCGATCTA |
| CBX3-gRNA4-S | TTCTTAACTCTCAGAAAGC |
| CBX3-gRNA4-AS | GCTTTCTGAGAGTTAAGAA |
| CBX3-gRNA5-S | AACCAAGAGGATTTGCCAG |
| CBX3-gRNA5-AS | CTGGCAAATCCTCTTGGTT |

| **shRNA sequences for human HOXA9 or SAFB** |
|---|
| hHOXA9_sh-**1296-**Fwd |
| sequence: |
| 5'CCGGACGCTTGACACTCACACTTTGCTCGAGCAAAGTGTGAGTGTCAAGCGTTTTTT-3' |
| hHOXA9_sh-**1296-**Rev |
| sequence: |
| 5'-AATTAAAAAACGCTTGACACTCACACTTTGCTCGAGCAAAGTGTGAGTGTCAAGCGT-3' |
| hHOXA9_sh-**1018-**Fwd |
| 5'-CCGGACGCTTGACACTCACACTTTGCTCGAGCAAAGTGTGAGTGTCAAGCGTTTTTT-3' |
| hHOXA9_sh-**1018-**Rev |
| 5'-AATTAAAAAACGCTTGACACTCACACTTTGCTCGAGCAAAGTGTGAGTGTCAAGCGT-3' |
| hHOXA9_sh-**1293-**Fwd |
| 5'-CCGGTGGTTCTCCTCCAGTTGATAGCTCGAGCTATCAACTGGAGGAGAACCATTTTT-3' |
| hHOXA9_sh-**1293-**Rev |
| 5'-AATTAAAAATGGTTCTCCTCCAGTTGATAGCTCGAGCTATCAACTGGAGGAGAACCA-3' |
| hSAFB_sh-**2100-**Fwd |
| Forward sequence: |
| 5'-CCGGCGGACTGTAGTAATGGATAAACTCGAGTTTATCCATTACTACAGTCCGTTTTT-3' |
| hSAFB_sh-**2100-**Rev |
| Reverse sequence: |
| 5'-AATTAAAAACGGACTGTAGTAATGGATAAACTCGAGTTTATCCATTACTACAGTCCG-3' |
| hSAFB_sh-**1888-**Fwd |
| Forward sequence: |
| 5'-CCGGGGTGGTAATCCTGACGAAATTCTCGAGAATTTCGTCAGGATTACCACCTTTTT-3' |
| hSAFB_sh-**1888-**Rev |
| Reverse sequence: |
| 5'-AATTAAAAAGGTGGTAATCCTGACGAAATTCTCGAGAATTTCGTCAGGATTACCACC-3' |
| hSAFB_sh-**4945-**Fwd |
| Forward sequence: |
| 5'-CCGGAGAGGACAAAGAAACTATAAACTCGAGTTTATAGTTTCTTTGTCCTCTTTTTT-3' |
| hSAFB_sh-**4945-**Rev |
| Reverse sequence: |
| 5'-AATTAAAAAAGAGGACAAAGAAACTATAAACTCGAGTTTATAGTTTCTTTGTCCTCT-3' |

| mRNA Primers | | |
|---|---|---|
| **Name** | | **Sequence** |
| hHOXA9-qPCR-F1 | | TACGTGGACTCGTTCCTGCT |
| hHOXA9-qPCR-R1 | | CGTCGCCTTGGACTGGAAG |
| hHOXA9-qPCR-F2 | | GTCCAAGGCGACGGTGTTT |
| hHOXA9-qPCR-R2 | | CCGACAGCGGTTCAGGTTTA |
| hHOXA9-qPCR-F3 | | CTGTCCCACGCTTGACACTC |
| hHOXA9-qPCR-R3 | | CTCCGCCGCTCTCATTCTC |
| hSAFB-qPCR-F1 | | CAACAAGAGCGTTTTGATGGAG |
| hSAFB-qPCR-R1 | | TGTTTCCCTCGGAGGTAATTTCA |
| hSAFB-qPCR-F2 | | TACCTCCGAGGGAAACAAGAA |
| hSAFB-qPCR-R2 | | CAGCCCGTTATCTTCCACACC |
| hSAFB-qPCR-F3 | | TCGCAGCAGTTGTGGTAGAAA |
| hSAFB-qPCR-R3 | | TGACAAAACCGTAACAGCGAG |
| hGAPDH_F-mRNA | | CCACATCGCTCAGACACCAT |
| hGAPDH_R-mRNA | | CCAGGCGCCCAATACG |
| hCEBPD_F--mRNA | | GGTG CCCGCTGCAGTTT |
| hCEBPD_R –mRNA | | CTCGCAGTTTAGTGGTGGTAAGTC |
| hCDKN1A_F1-mRNA | | TGTCCGTCAGAACCCATGC |
| hCDKN1A_R1-mRNA | | AAAGTCGAAGTTCCATCGCTC |
| hCDKN1A_F2 -mRNA | | CGATGGAACTTCGACTTTGTCA |
| hCDKN1A_R2-mRNA | | GCACAAGGGTACAAGACAGTG |
| hNOTCH1-F1-mRNA | | GAGGCGTGGCAGACTATGC |
| hNOTCH1-R1-mRNA | | CTTGTACTCCGTCAGCGTGA |
| hNOTCH1-F2-mRNA | | TGGACCAGATTGGGGAGTTC |
| hNOTCH1-R2-mRNA | | GCACACTCGTCTGTGTTGAC |
| ABLIM3-mRNA-F1 | | TCC GCG TGC ACA ACA ACC ACT T |
| ABLIM3-mRNA-R1 | | GCT GTC ACA GCG GGT GCC ATA G |
| ABLIM3-mRNA-F2 | | CAA ATC TGC CTC CCT GCC TGC C |
| ABLIM3-mRNA-R2 | | GCA TGC CCC AGT TGA CTG CGT T |
| S100A12-mRNA-F 2 | | ACA AAG GAG CTT GCA AAC ACC A |
| S100A12-mRNA-R 2 | | GCC TTC AGC GCA ATG GCT ACC A |
| S100A12-mRNA-F 1 | | CAG TGC CCT TCA CCA CTG CTG G |
| S100A12-mRNA-R 1 | | TGC CCC TTC CGA ACT GAG TAT TGG T |
| S100A9-mRNA- FWD 1 | | GGG GGA ATT CAA AGA GCT GGT GCG |
| S100A9-mRNA- REV 1 | | CGA AGC TCA GCT GCT TGT CTG CA |
| S100A9-mRNA-FWD 2 | | ACA CTC TGT GTG GCT CCT CGG C |
| S100A9-mRNA- REV 2 | | TGT TGC GTT CCA GCT GCG ACA T |
| S100A8-mRNA- FWD 1 | | TGC CTC TCA GCC CTG CAT GTC T |
| S100A8-mRNA-REV 1 | | ACA TGA TGC CCA CGG ACT TGC C |
| S100A8-mRNA-FWD 2 | | AGG GTG CAG ACG TCT GGT TCA A |
| S100A8-mRNA- REV 2 | | GGC TGC CAC GCC CAT CTT TAT C |
| BCL2A1-mRNA- FWD 1 | | ACG ACA GCA AAT TGC CCC GGA T |
| BCL2A1-mRNA- REV 1 | | TTC CCA GCC TCC GTT TTG CCT T |
| BCL2A1-mRNA-FWD2 | | AAC GGA GGC TGG GAA AAT GGC T |
| BCL2A1-mRNA-REV2 | | TGG AGT GTC CTT TCT GGT CAA CAG T |
| NCF1-mRNA-F1 | | TAC ACT GCT GTG GAG GGG GAC G |
| NCF1-mRNA-R1 | | TTC CTG ATG ACC CAC CAG CCG T |
| NCF1-mRNA-F2 | | GAC GGC TGG TGG GTC ATC AGG A |
| NCF1-mRNA-R2 | | TGG GAC ACG TCT TGC CCT GAC T |
| GATAD2A-mRNA-F1 | | CGT CAA CAT CCC ACA GCC CAC C |
| GATAD2A-mRNA-R1 | | TCG GCA GAG GTG ACC ACA GAG G |
| GATAD2A-mRNA-F2 | | ACA AAA GGA AGC CAC CGC CCA G |
| GATAD2A-mRNA-R2 | | GGA GTG ATG GCT TGC CAG CAG G |
| MTA2-mRNA-F1 | | ACA ACC CTC TCA CAG ACC GGC A |
| MTA2-mRNA-R1 | | GGC TGC CGA ATG GAG CTG CTA C |
| MTA2-mRNA-F2 | | CAT TCG GCA GCC AAG CTT GCA C |
| MTA2-mRNA-R2 | | AGC CAG GTC GTA GCC GTT CCT T |
| CBX3-mRNA-F1 | | TGC TGC TGA CAA ACC AAG AGG A |
| CBX3-mRNA-R1 | | CCA CTG CTG TCT GTG GCA CCA A |
| CBX3-mRNA-F2 | | GGC AGA GCC TGA AGA ATT TGT CGT GG |
| CBX3-mRNA-R2 | | TCA GGT TCC CAA GTA TTG TCA GCA |

| Antibodies | | | |
|---|---|---|---|
| **Antibody** | **Species** | **Catalogue No** | **Company** |
| Anti-SAFB clone 6F7 | Mouse monoclonal IgG | 05-588 | Millipore |
| Anti-HOXA9 | Rabbit polyclonal | HPA061982 | ATLAS Anitobodies |
| CRISPR/Cas9 | Mouse monoclonal IgG1k | C15200203 | diagenode |
| MTA2 Antibody | Rabbit polyclonal | A300-395A | BETHYL |
| p66alpha/GATAD2A Antibody | Rabbit polyclonal | A302-356A | BETHYL |
| CBX3 Antibody | Rabbit polyclonal | A300-984A | BETHYL |
| Histone H3 (1B1B2) | Mouse monoclonal IgG | 14269 | Cell Signaling |
| Anti-Histone H3 (tri methyl K9) antibody | Rabbit polyclonal | ab8898 | Abcam |
| Di-Methyl-Hisotne H3 (Lys9) (D85B4) | Rabbit monoclonal | 4658 | Cell Signaling |
| Anti-Histone H3 (di methyl K9) antibody [mAbcam 1220] | Mouse monoclonal | ab1220 | Abcam |
| Tri-Methyl-Hisotne H3 (Lys27) (C36B11) | Rabbit monoclonal | 9733 | Cell Signaling |
| Tri-Methyl-Hisotne H3 (Lys4) (C42D8) | Rabbit monoclonal | 9751 | Cell Signaling |
| CEBPd (C6) | Mouse monoclonal | sc-365546 | SanatCruz Biotechnology Inc. |
| IgG control Rabbit polyclonal antibody | Rabbit | 30000-0-AP | Proteintech |
| Peroxidase conjugated Goat-Anti-Rabbit IgG, Fc Fragment specific | Goat-Anti-Rabbit | 111-035-046 | Jackson ImmunoResearch |
| Peroxidase conjugated Goat-Anti-Mouse IgG, Light Chain specific | Goat-Anti-Mouse | 115-035-174 | Jackson ImmunoResearch |
| Peroxidase conjugated Monoclonal Mouse Anti-Rabbit IgG IgG, Light Chain specific | Mouse Anti Rabbit | 211-032-171 | Jackson ImmunoResearch |

| Inhibitors | | | | | | | | |
|---|---|---|---|---|---|---|---|---|
| **Inhibitor** | **Catalog No.** | | **Company** | **CAS No.** | **Formula** | **Molecular Weight** | **Dissolved in DMSO** | |
| **Panobinostat (LBH589)** | S1030 | Synonyms: NVP-LBH589 | Selleck | 404950-80-7 | $C_{21}H_{23}N_3O_2$ | 349.43 | 10mM | Panobinostat (LBH589, NVP-LBH589) is a novel broad-spectrum **HDAC** inhibitor with **IC50** of 5 nM in a cell-free assay |
| **Chaetocin** | HY-N2019 | | MedChemExpress (MCE) | 28097-03-2 | $C_{30}H_{28}N_6O_6S_4$ | 696.84 | 10mM | Chaetocin is a specific inhibitor of the histone methyltransferase (**HMT**) SU(VAR)3-9 with an **IC**$_{50}$ of 0.6 μM for SU(VAR)3-9 |
